# Tissue engineered model of hepatic breast cancer micrometastasis shows host-dependent colonization patterns and drug responses

**DOI:** 10.1101/2020.01.08.898163

**Authors:** Anna Guller, Vlada Rozova, Inga Kuschnerus, Zahra Khabir, Annemarie Nadort, Alfonso Garcia-Bennett, Liuen Liang, Yi Qian, Ewa M. Goldys, Andrei V. Zvyagin

**Author notes:** Corresponding authors: Anna Guller, Ewa Goldys.

## Abstract

Early stages of colonization of distant organs by metastatic cancer cells (micrometastasis) remain almost inaccessible to study due to lack of relevant experimental approaches. Here, we show the first 3D tissue engineered model of hepatic micrometastasis of triple negative breast cancer (TNBC). It reproduces characteristic histopathological features of the disease and reveals that metastatic TNBC cells colonize liver parenchymal and stromal extracellular matrix with different speed and by different strategies. These engineered tumors induce the angiogenic switch when grafted in vivo, confirming their metastatic-specific behaviour. Furthermore, we proved feasibility and biological relevance of our model for drug and nanoparticle testing and found a down-regulatory effect of the liver microenvironment of the sensitivity of TNBC cells to chemotherapeutic drug doxorubicin in free and nanoformulated forms. The convenient and affordable methodology established here can be translated to other types of metastatic tumors for basic cancer biology research and adapted for high-throughput assays.

## INTRODUCTION

Metastatic cascade includes the dissemination of cancer cells from the primary tumor to the distant organs mainly through the blood and lymphatic vessels, their escape from the vasculature (extravasation), homing in a secondary organ and its colonization. The initial metastatic homing is extremely rare event, as only 0.01% of circulating cancer cells can attach in the secondary organ microenvironment and start a new colony ^1^. If not destroyed by immune system, this colony may further develop into avascular micrometastases and stay dormant or, through transient acceleration of tumor-induced blood vessels growth in the vicinity of the cancer cell colony known as angiogenic switch ^2^, progress to massive blood-perfused macrometastases ^3, 4^.

In contrast to macrometastases, the early stages of metastatic cascade (from homing to angiogenic switch) are difficult to detect in natural conditions and to study in the lab. Indeed, spontaneous metastatic spreading is widely addressed in the literature with a number of experimental models of the induction or distant metastases, mainly by direct injections of cancer cells into vasculature followed by observations of the macroscale metastatic lesions forming in the distant organs. At the same time, the studies of the latent post-extravasation events like cancer cells homing and metastatic colonization of secondary organs as well as the “mysterious” specific organotropism of the certain types of tumors, known by Paget’s “seed and soil” metaphor, are rare ^4^ and the most experimentally challenging because of lack of reliable methodologies ^5^.

Hepatic metastases of triple-negative breast cancer (TNBC) is a particularly bitter example of this knowledge gap ^6^. TNBC is an aggressive HER2^-^, estrogen and progesterone receptor-negative mammary carcinoma ^6-8^. Compared with other breast cancer types, TNBC carries ∼2.5 times higher risk of distant metastases within 5 years after diagnosis, resulting in much poorer prognosis than other types of breast carcinomas with the excessive death rates of 70% versus 44%, respectively ^9^. Lack of targeted therapy and limited efficiency of conventional treatment (mainly chemotherapy with cytostatic drugs) options particularly contribute to the low survival of the patients with metastatic TNBC ^10^. Up to one half of TNBC metastases occur in the liver ^7, 11, 12^. The causes of such liver preference of the TNBC metastases are virtually unknown. As the role of blood inflow is rather comparable to other breast carcinomas, the local tissue-specific mechanisms are hypothesized to be a key factor ^5, 13^. In particular, special organization of hepatic vasculature, such as discontinuous basement membranes in sinusoids, allows extravasation of metastatic cancer cells to occur first in the narrow (∼1-2 μm) gap between endothelial cells and hepatocytes, known as Disse’s space ^14-16^. It forms the major part of ECM framework of hepatic parenchyma and contains blood plasma, collagens types I, III, V, VI and VII, as well as fibronectin and tenascin ^17^. Therefore, hepatic extracellular matrix (ECM) makes the first homing space for the incoming cancer cells ^14, 18, 19^ and becomes the potential starting point, the operational environment and the regulator of metastatic colonization of the organ ^20, 21^. Therefore, the biologically relevant and reproducible models of TNBC hepatic metastases must reflect the cancer cells - liver ECM interactions and include at least two components: the representative TNBC cells and the biocompatible material playing the role of the liver ECM. We believe that these models are essential for the advancement in understanding, prevention and treatment of the TNBC metastases to the liver.

Stable linear cells are a reasonable choice to ensure biological accuracy and reproducibility of the models. Among TNBC-related lines, MDA-MB-231 is recognized as a triple-negative, mesenchymal-like invasive cell phenotype in vitro, which is also tumorigenic in vivo ^22^ and commonly favoured for simulations of the metastatic-related features of TNBC. However, two-dimensional (2D) monolayer cell cultures in vitro lack biological accuracy due to almost total absence of cell-ECM interactions, artificial stiffness and geometry of the plastic cultural substrate and abnormal mass-transport, which are critical for cancer biology research and drug testing ^23, 24^. The most common three-dimensional (3D) in vitro cultures such as multicellular spheroids of TNBC cells are considered as a model of small solid avascular microtumours ^25, 26^, where cell-ECM interactions are overrun by cell-cell communications. Considering the liver-specific conditions discussed above, this rather reflects the structure of the internal parts of the massive tumors than the early metastatic colonies. In order to “add” cell-matrix interactions to 3D culture systems, cells or spheroids are conventionally cultured in or on artificial gel-like materials. However, this approach does not reproduce realistic organ-specific tissue mechanics ^27^, while mechanotransduction is critically important in cancer biology ^28^.

An alternative methodology of tumor tissue engineering ^29, 30^ relies on creation of 3D tissue engineering constructs (TECs) combining cells of interest and *solid* scaffolds. These scaffolds include porous and fibrous biomaterials engineered (casted, moulded, 3D-printed, electrospun etc.) on the base of various polymers or prepared by *decellularization* of natural tissues or cells ^29, 31^. Decellularization (DCL)^32, 33^ is removal of the cells from the tissues or cellular sheets with preservation of the ECM. Decellularized tissues are among the most clinically successful biomaterials in regenerative medicine because they preserve three essential organ-specific features of ECM: the composition, the architecture and the biomechanical properties ^34^. It is generally held that the best quality of DCL is achieved by perfusion of the whole organs via natural vasculature (WO-DCL) in a bioreactor ^33^. The procedure requires extremely careful cannulation of the major blood vessels in a live anesthetized animal; and therefore, it is almost non-scalable, low-reproducible and non-applicable in pharmacological research. This promotes the trend of reduction of the biological accuracy of the scaffolds as ECM replicas if favour of their reproducibility by using artificial materials. For example, TNBC metastases to the bone was mimicked in a tissue engineering model using of MDA-MB-231 cells seeded on polycaprolactone ^35^ and silk ^36^ scaffolds and tested in murine hosts. Some studies demonstrate merging DCL-based and polymer-based scaffolding. Using polycaprolactone scaffolds coated with ECM extracts of decellularized liver and lungs, Aguado et al. show that organ-specific ECM stimulates colonization of pre-metastatic niche by TNBC cells ^37^. However, to the best of our knowledge, non-transformed (i.e., not solubilized, lyophilized or chemically cross-linked) decellularized organ-specific matrix was applied in only two short-term (up to 7-9 days in vitro) model studies addressing growth and metastatic invasion of breast cancer cells into the lung ECM ^38, 39^, but it has been never used in modelling of hepatic metastases of TNBC.

In current study we present a new tissue engineering model or early metastasis of TNBC to the liver, which includes MDA-MB-231 cells and organ-specific ECM of macroscale (8-10 mm^3^) chick embryo liver scaffolds prepared by an original immersion whole-organ decellularization (iWO-DCL) procedure. In contrast to standard perfusion WO-DCL of single ^40^ or multiple ^41^ organs, our iWO-DCL method does not rely on precise surgical procedures and blood vessels cannulation, but can be efficiently applied to various *whole organs* of small vertebrates. Considering cross-species similarity of ECM ^32^ of the same organs, we performed the whole study using well-known, reproducible and affordable chick embryo experimental platform in order to facilitate further use of our methodology in a wide range of cancer biology and drug development research and industrial applications.

Using our model, we revealed strikingly different patterns in attachment, migration, growth and matrix remodelling behaviour of metastatic TNBC cells in parenchymal and stromal ECM of the liver, respectively, as well as reversible adaptations of the cellular epithelial- and mesenchymal-like morphotypes at different stages of colonization. This has potential clinical relevance indicating the need for a local regional-specific approach to diagnostics and treatment of TNBC metastatic lesions. The ability of TECs to initiate growth of new blood vessels in the chick embryo host tissue in vivo confirmed the cancer behaviour relevance of the model. Finally, in our study we found significant differences in growth dynamics of TNBC cells on 2D culture plastic and in 3D hepatic ECM. This corresponded with the dramatic reduction of therapeutic efficiency (as cytotoxicity towards the cancer cells) and cellular uptake of free Doxorubicin hydrochloride (Dox), a cytostatic drug widely used in chemotherapy of primary and metastatic TNBC, and original anionic surfactant template mesoporous silica nanoparticles (AMS-6), loaded with Dox (AMS-6-Dox) in our 3D engineered model, in comparison with matching 2D in vitro cultures of MDA-MB-231 cells. These findings show the effect of liver-specific ECM of metastatic TNBC cells drug sensitivity and demonstrate feasibility of our new organ-, disease- and stage-specific TNBC biology model as a more biologically accurate testbed for drug development and nanomedicine, potentially applicable for high-throughput assays.

## RESULTS

### Study design

Our study is schematically illustrated in Figure 1. We used chick embryos incubated until the embryonic day 18 (ED18) when all major organs have formed ^42^, while pain sensitivity is not yet fully developed. The iWO-DCL of the native chick embryo liver, described in the Methods section and illustrated in Supplementary Figures S2-S4, was used to produce acellular organ-specific scaffolds (AOSS) as 8-10 mm^3^/2-3 mm side size fragments of decellularized chick embryo liver. Next, TECs were developed via seeding of AOSS with MDA-MB-231 cells. The cells grew on and within the scaffolds in a static *in vitro* culture for up to 4 weeks allowing us to characterize the cell invasion patterns, examine drug response and validate *in vivo* the cancer-specific behaviour after implantation in chick embryos. The TECs were sampled for histological and other analyses at various time points during the *in vitro* culture. Matching 2D *in vitro* cultures of the same cells were studied in parallel with the TECs. The TECs with well-developed cell colonies (after 3 weeks of *in vitro* culturing) were used for evaluation of cytotoxicity and cellular uptake of doxorubicin (Dox) as well as Dox-loaded mesoporous silica nanoparticles (Anionic Mesoporous Silica-6, AMS-6-Dox ^43^). The ability of the engineered tumors to induce the “angiogenic switch”, or growth of blood vessels in the host tissue, was examined by chick embryo chorioallantoic membrane (CAM) assay via grafting of the TECs ^44^. This effect is known to be a critical cancer hallmark ^45^ especially significant for metastatic progression ^2-4^.

**Figure 1.**
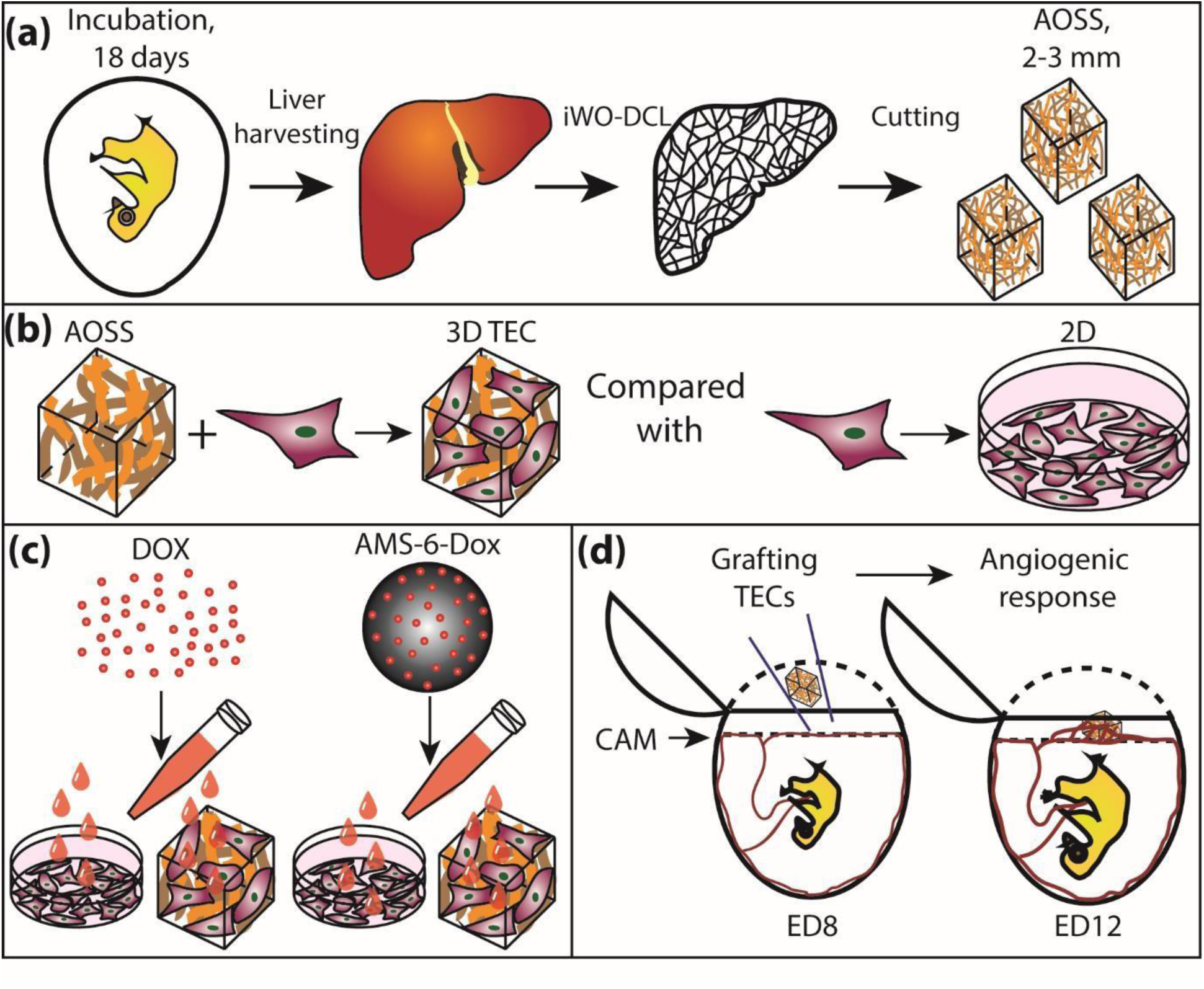
Schematic illustration of study design. (a) Preparation of the liver AOSS: harvesting liver from ED18 chick embryos, iWO-DCL of the livers, and preparation of the AOSS by cutting decellularized livers into small fragments; (b) Preparation of 3D TECs by seeding of the liver AOSS with MDA-MB-231 cells; analysis of the TNBC colonization patterns and cellular geometries in TECs and comparison of cell population dynamics in TECs with conventional 2D cell cultures; (c) Comparison of the effect of molecular Dox and nanoparticleformulated AMS-6-Dox in a 2D culture of TNBC cells and in 3D TECs; (d) Evaluation of angiogenesis induced by TECs grafted on CAM. Abbreviations used in the figure: AOSS – acellular organ-specific scaffold; DCL – decellularization; ED – embryonic day (age of the chick embryo); iWO-DCL – immersion-agitation-assisted whole-organ DCL; Dox – doxorubicin; AMS-6-Dox – mesoporous silica nanoparticles loaded with 20% doxorubicin; TEC – tissue engineering construct; 2D and 3D – two-dimensional TNBC cell cultures and 3D TECs.

### Histological evaluation of cell growth patterns in TECs

The details of the native structure of chick embryo liver and the effects of applied iWO-DCL procedure are described in SI and Figures S1-S5. Briefly, after removal of cells, the liver ECM was found to be well preserved and had the composition and spatial structure similar to that of the liver of other vertebrates decellularized by perfusion WO-DCL ^46-48^. Two histoanatomical compartments were observable in the decellularized chick embryo liver and randomly appeared in AOSS (Figure S3); their origin and composition are schematically illustrated in Figures S4 and S5. The first compartment contained sponge-like structures corresponding to the former hepatic parenchyma and formed mainly by the residuals of the Disse’s space elements (Figure 2 (a)). The second compartment was formed by denser structures relatively enriched with fibrillar collagen mostly corresponding to the former walls of central veins, portal triads, and interlobular connective tissue sheaths and capsular connective tissue (Figure 2 (b)). The matrix appeared densified and located at the edges of the scaffolds where the morphological features were less discernible was also considered as a part of the second compartment in further quantitative analysis without regard of its’ actual origin. Below, we term the first compartment as “parenchymal” and the second as “stromal” for simplicity.

**Figure 2.**
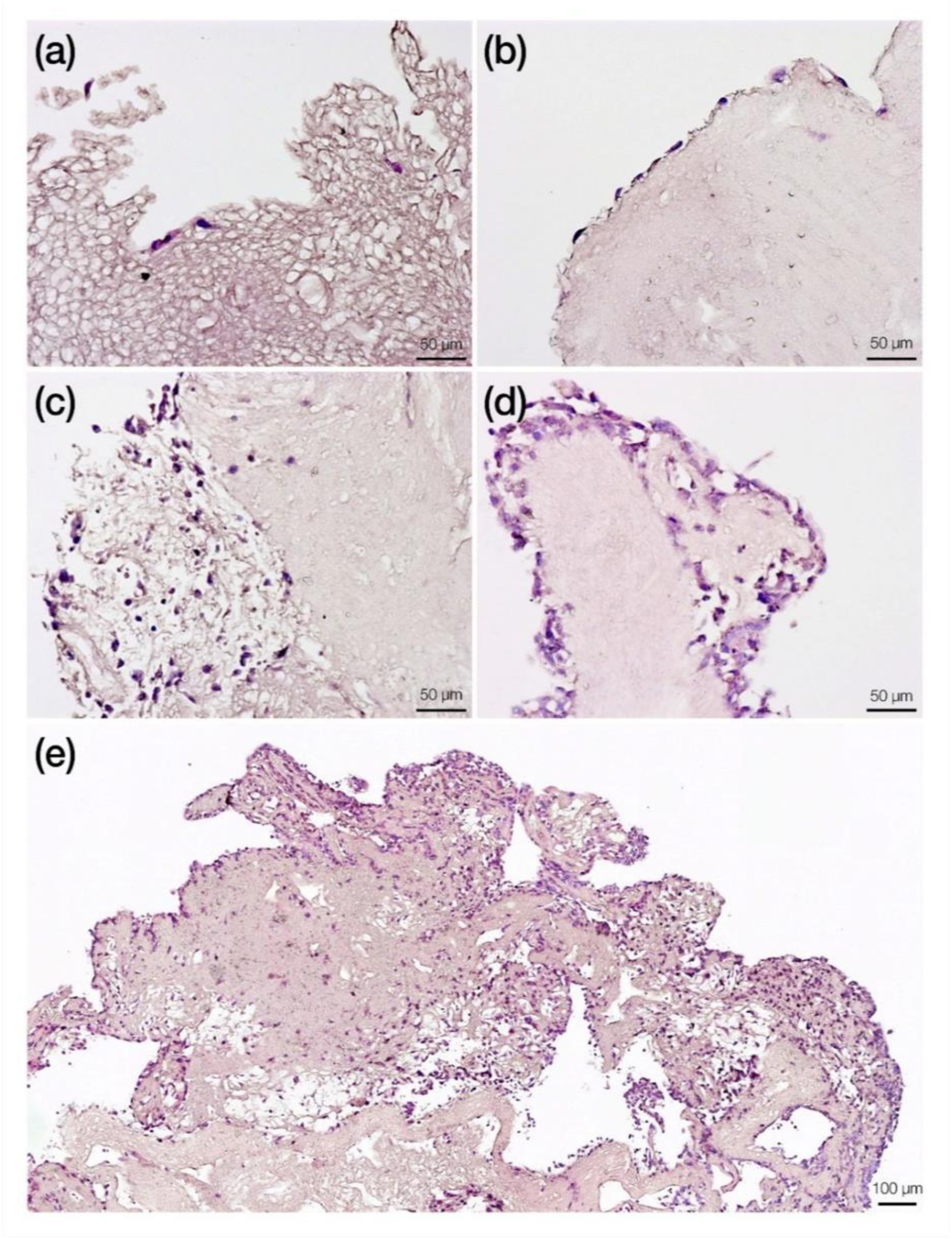
Representative histology images of cellular colonization of the liver AOSS. (a) The parenchymal compartment on Day 5. Sparse individual cells are attached to the loose spongy-like ECM on the outer border of the TEC. (b) Formation of a discontinuous lining on the dense ECM of the stromal compartment (Day 5). (c) Colonization of the parenchymal compartment (the fragment on the left) by individual cells and small multicellular clusters showing predominant single-cell invasion (Day 13). (d) Continuous multi-row cell lining on the stromal ECM with minimal invasion (Day 13). (e) Remodelling of the ECM of stromal origin and massive diffuse colonization of the whole scaffold (Day 28).

The histological study and digital image analysis of TECs over 4 weeks of *in vitro* culturing revealed distinctive patterns of initial cellular attachment and subsequent metastatic colonization in the parenchymal and stromal ECM compartments (Figure 2 and Table S1). Moreover, cells exhibited phenotypical adaptations reflected in their morphology, specifically the circularity (*c*, ratio of cell area to its squared perimeter). Two subpopulations with varying cell circularity could be observed, further referred to as “epithelioid” (*c ∼* 0.85) and “mesenchymal-like” (*c ∼* 0.7) as shown in Figure S6.

During *Week 1* of the *in vitro* TEC culture, the number of cells attached to the ECM of the stromal compartment was 2.8 times higher than to the parenchymal ECM (Figure 2 (a, b)). Most cells observed in the stromal compartment [63% (±10%)] had a mesenchymal-like elongated appearance, while in the parenchymal compartment a slight majority of cells had more circular epithelioid morphology [59% (±16%)], and the rest appeared mesenchymal-like. Notably, at this time point, most cells in the in the stromal compartment formed single-row linings at the outer surfaces of the ECM, while the cells in the parenchymal compartment were mainly distanced from each other.

At the end of *Week 2*, in the parenchymal compartment, individual cell colonization was predominantly [75% (±23%)] observed, while the remaining cells formed small multicellular clusters and single-cell surface linings over the AOSS surface (Figure 2 (c)). These individual cells mainly had the epithelioid morphology [76% (±7%)], and the rest appeared mesenchymal-like. Cells in the stromal compartment mostly aggregated in multicellular clusters, both on the scaffold surfaces (as double- or triple-row linings or islands) and in the depth of the matrix, and only 17% (±21%) of cells in the stromal compartment remained single (Figure 2 (d)). The invasion of the stromal compartment by the cell clusters occurred primarily along voids and clefts of the ECM. These clusters invaded the ECM to a depth (defined as a distance from the AOSS surface) estimated as 70 µm (± 24 µm), in contrast to the deeper (134 µm (±54 µm)) invasion by individual cancer cells in the parenchymal compartments.

During *Weeks 3* and *4*, a notable sponge-like remodelling of the ECM was observed in the stromal compartments, resulting in a blurred distinction between the ECM of the parenchymal and stromal origins. The invasion of cancer cells progressed over this period, reaching up to 800 µm depth from the AOSS surface regardless of the matrix origin (Figure 2 (e)).

By the end of *Week 4*, the average cellular density across whole TECs was almost the same as that in the parenchymal compartment on *Week 2* (Table S1), and, approximately 70% (±.16%) of single cells in all compartments retained their epithelioid morphology.

### Growth dynamics in 3D TECs and 2D cultures of MDA-MB-231 cells

A comparative analysis of the cell growth in 2D and 3D TECs *in vitro* cultures was carried out using the MTT assay (Figure 3 and Table S2). The number of viable cells successfully attached to the surface of AOSSes on Day 1 was found to be ∼15% of the live cells’ population in the matching 2D culture. The cells in 2D cultures and in TECs presented strikingly different growth behaviour. As it is shown in Figure 3 (a), cell density increased until Day 21 in both cultures. During *Week 1*, the growth observed in the 2D cultures was much faster than in the TEC counterparts (Figure 3 (b)). In the following 2 weeks, the weekly cell count increased similarly in the 2D cultures and 3D TECs due to dramatic deceleration of growth rate in monolayers and slow but steady proliferation of cells in TECs. In final *Week 4*, the growth rates in both types of cultures decreased in comparison to that of *Week 3*. This decrease was less pronounced in the TECs. Between Days 21 and 28, the cell density in 2D cultures decreased by approximately 19% (±10%) to 2.4×106 cells per sample; and in the TECs – by 8% (±36%) to 4.4×105 cells per sample. Differences between the viable cell counts in 2D and TEC cultures were statistically significant at each time point (p <0.001).

**Figure 3.**
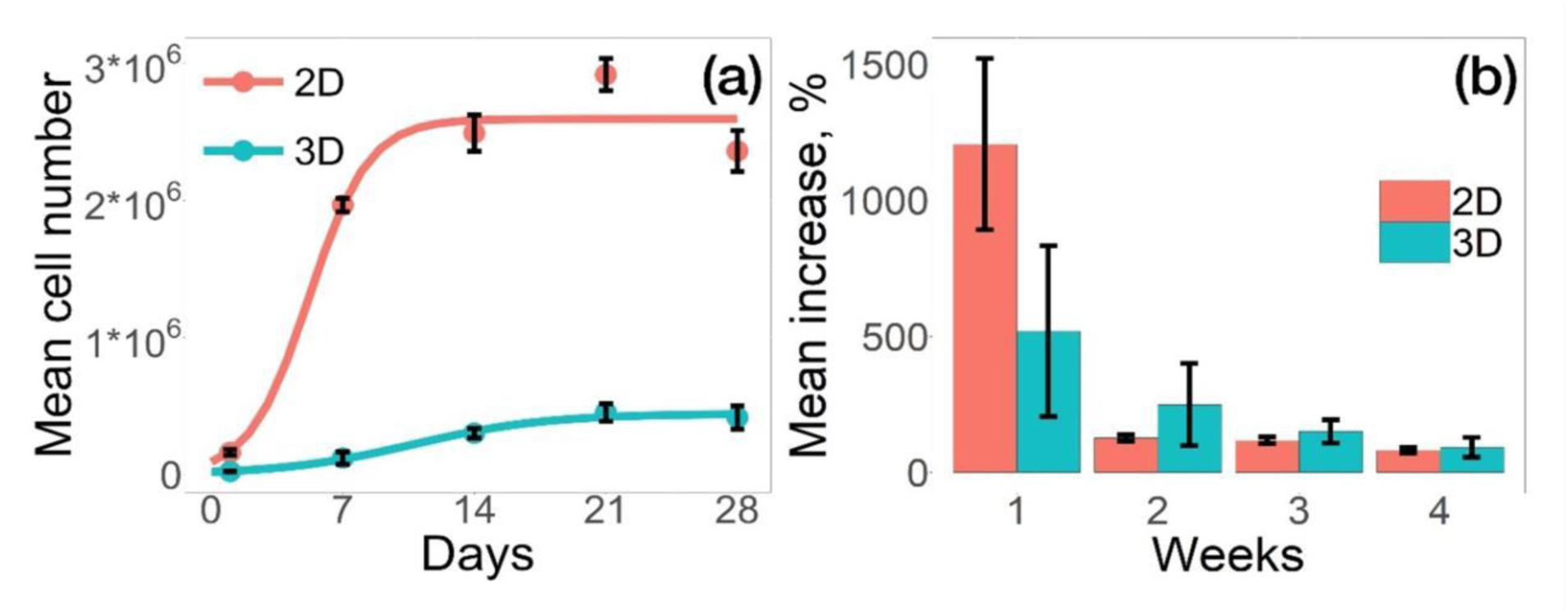
Cell growth dynamics of MDA-MB-231 cells in 2D and 3D in vitro cultures evaluated using MTT assay. (a) Average number of cells in both cultures. Solid curves represent the lines of best fit given by Equation 3; error bars indicate 95% CIs. (b) Average weekly increases of the cell numbers in 2D and 3D cultures. Error bars indicate standard deviation.

The observed cell population dynamics in the matching 2D and 3D *in vitro* cultures of MDA-MB-231 cells was found to closely follow a general logistic growth model (see Equation 3, Table S2). Both the maximum cell number (*C*_*max*_) and cell population growth rate (*d*) in 3D TECs were lower compared to the corresponding values in the 2D cell culture (*C*_*max*_ = 4.4×10^5^ cells, *d* = 0.28 days ^-1^ in 3D vs. *C*_*max*_ = 2.4×10^6^ cells, *d* = 0.63 days ^-1^ in 2D).

### Cytotoxicity and cellular uptake of Dox and AMS-6-Dox in 3D TECs and 2D in vitro cultures of MDA-MB-231 cells

Nanoformulated AMS-6-Dox was prepared and carefully characterized, as described in the Methods, and the results are shown in Figures S7-S13. The results of MTT viability assays performed in 2D and 3D TEC cultures incubated for 36 h in the presence of molecular and nanoformulated Dox are shown in Figure 4 and in Figure S14. We found that pristine AMS-6 nanoparticles were not cytotoxic in both 3D TECs and 2D cultures at a concentration range 0 – 250 µg/mL, which is consistent with our previous studies ^49^.

**Figure 4.**
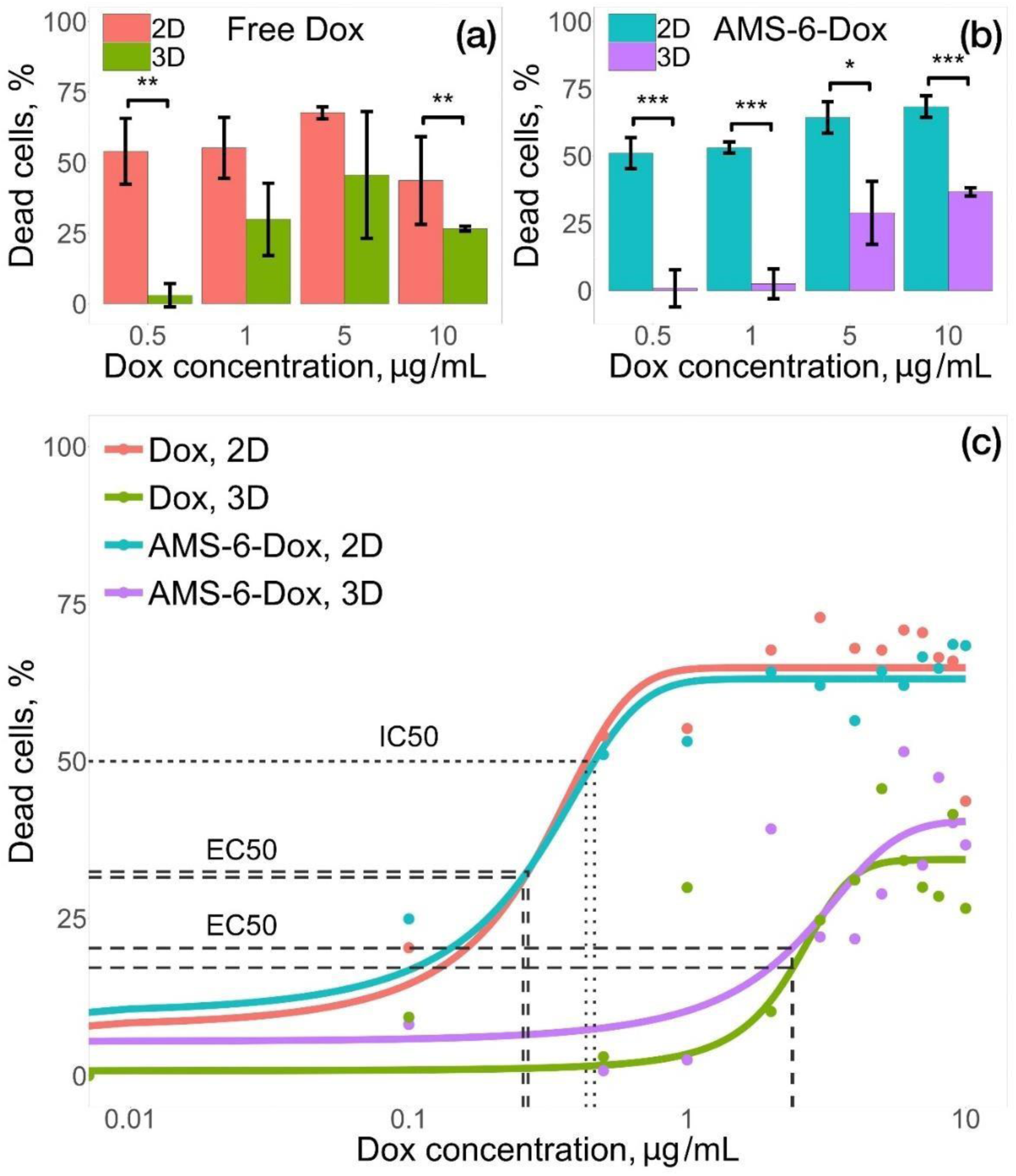
Results of MTT test of cytotoxicity of free and nanoformulated Dox in 2D cultures of MDA-MB-231 cells and 3D TECs. (a) The effect of free Dox on MDA-MB-231 cells in 2D and 3D in vitro cultures. (b) Effect of AMS-6Dox on MDA-MB-231 cells in 2D and 3D in vitro cultures, exposure time, 36 h. Error bars indicate 95% CIs; statistically significant difference at *p < 0.05, **p < 0.01, ***p < 0.001 by Mann-Whitney U test. (c) Dose-response curves of free Dox and AMS-6-Dox in 2D and 3D in vitro cultures. Solid lines represent the lines of best fit given by Equation 5. (See Tables S3 and S4 for fit parameters). Note that in the 3D culture, the IC50 could not be calculated within the studied concentration range (up to 10 µg/mL).

The cells in 3D TECs appeared much more resilient to both molecular and nanoformulated Dox in comparison with cells in 2D cultures (Figure 4 (a, b)). The dose-response relationship in each group was described, using a sigmoid function (see Methods for details, Equation 5). A half maximum effective concentration, EC_50_, in 3D TECs was found to be approximately 9 times higher than in 2D cultures for both formulations (Tables S4, S5). Moreover, while in 2D culture, the half maximum inhibitory concentration, IC_50_, was found to be 0.43 µg/mL and 0.46 µg/mL for free and nanoformulated Dox, respectively, in 3D TECs, 50% cell death threshold was not reached, indicating a markedly worse efficacy of both formulations in TECs compared to conventional 2D cultures (Figure 4 (c)).

Our data also shows the effect of Dox formulation (free or nanoformulated) on the cellular response in both 2D and TEC cultures (Figure 4 (a, b), Figure S15). Nanoformulated Dox was generally more cytotoxic at higher Dox concentrations (> 5 µg/mL), while the efficacy of free Dox was reduced at the higher Dox concentrations in both 2D and 3D cultures. At a maximum studied Dox concentration of 10 µg/mL the difference in mean viability between 3D and 2D cultures was 17.1% and 31.7% for Dox and AMS-6-Dox, respectively (see Figure 4 (a, b)).

The results of evaluation of cell and tissue uptake of Dox and AMS-6-Dox in 2D and 3D TEC cultures are shown in Figure 5. In 2D cultures, the fluorescent signals of free Dox and AMD-6-Dox nanoparticles were visible in the cellular nuclei (Figure 5 (a – h)). In contrast, in 3D TECs, fluorescence signals of free and nanoformulated Dox were distributed between the cell nuclei, cell cytoplasm and ECM of TECs. Free Dox was noticeably accumulated not only in the cell nuclei, but also throughout the TECs’ ECM, while fluorescence from AMS-6-Dox in 3D TECs was mainly detected in the nuclei of cells located at the TEC surface and only diffuse DAPI staining and Dox-positive cell debris was observed at a depth of > 50 µm in TEC, indicating destroyed nuclei and destroyed tumor cells, respectively (Figure 5 (i - p)). Taken together, the MTT test and imaging results show that the distribution and cytotoxic effects of free and nanoformulated Dox were influenced by the liver-specific ECM microenvironment, and the method of Dox delivery to 3D TECs modulated its local therapeutic activity.

**Figure 5.**
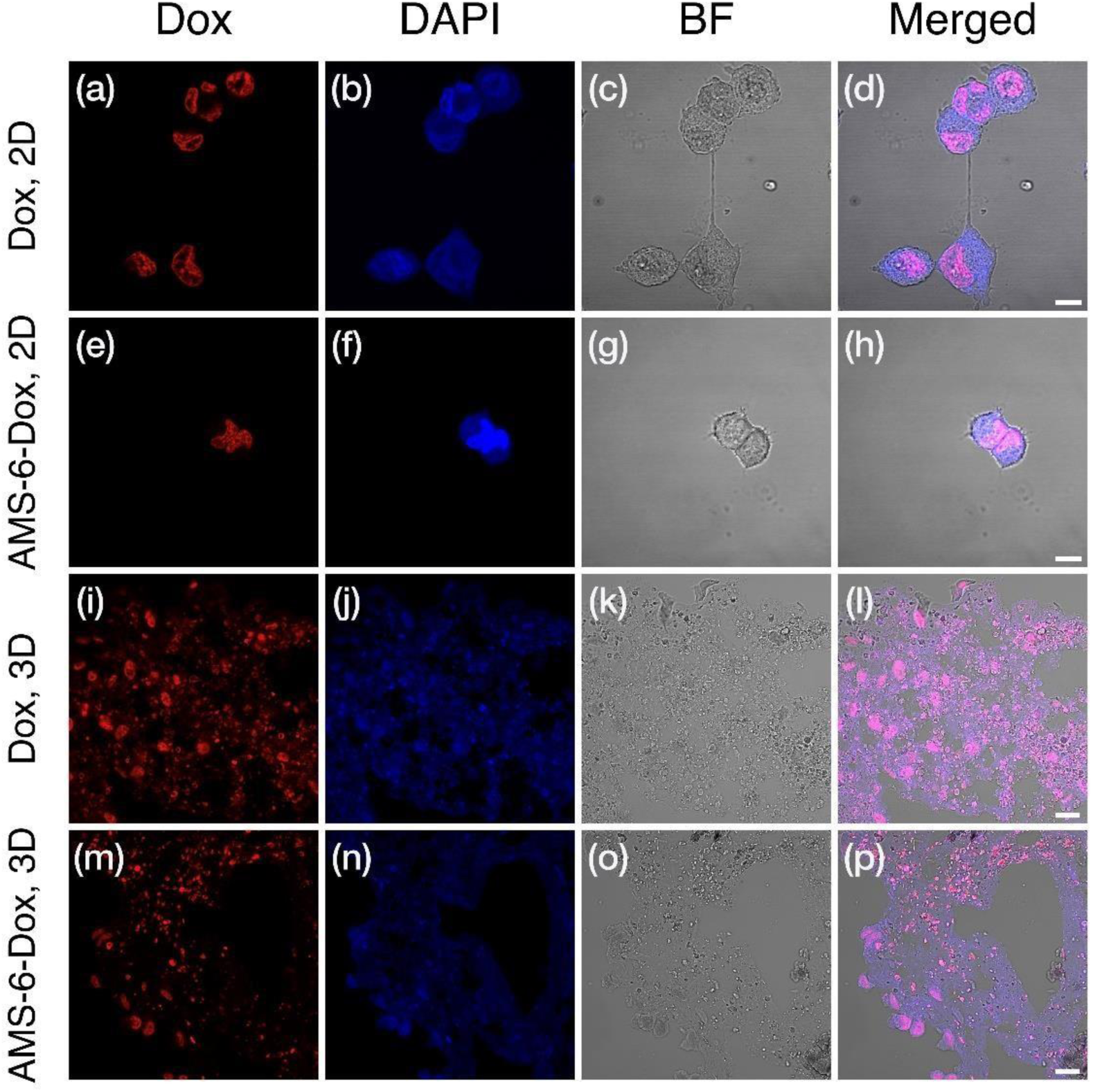
Confocal microscopy images of MDA-MB-231 cells incubated for 24 h with (a – d) Dox in 2D; (e – h) AMS-6-Dox in 2D; (i – l) Dox in 3D; and (m – p) AMS-6-Dox in 3D TECs. Intrinsic Dox fluorescence was detected in the red channel (Dox), while DAPI fluorescence (blue channel, DAPI) was used for staining of cell nuclei. Control bright field (BF) images were acquired to visualize the tissue structures, and merged images highlight the colocalization of Dox and DAPI signals. Note the absence of preserved cell nuclei in the depth of TEC treated with AMS-6-Dox nanoparticles (n), in comparison with TECs treated with free Dox (j). Dox concentration, 10 μg/mL. Scale bars, 10 μm (Dox, 2D; AMS-6-Dox, 2D) and 20 μm (Dox, 3D; AMS-6-Dox, 3D).

### Angiogenic potential of TECs and liver AOSSes in vivo

The angiogenic potential of the TECs and liver AOSSes in vivo was examined using the chick embryo CAM model. We found that grafting of CAM with 3D TECs, liver AOSSes and suspensions of MDA-MB-231 cells induced different changes of natural angiogenesis in CAM occurring between *ED8* and *ED12* and resulting in specific architecture of blood vessel trees in the studied groups. The convergence on the host blood vessels towards the graft was clearly visible in CAMs grafted with AOSSes and in CAMs grafted with TECs, but it was not obvious in other groups (Figure S19). The cellular xenografts initiated spatially irregular branching of small diameter vessels, while in the *Intact control, ED12* parallel large blood vessels were observed in combination with regularly arranged capillary meshes (Figure S19).

The TECs and cell xenografts induced statistically significant increase of branch length density in CAM blood vessels, in comparison to the intact control, while grafting of the AOSSes did not result in this angiogenic switch (Figure 6 (a-f) and Figure S18). In contrast to other types of xenografts, TECs also induced intensive branching of blood vessels (Figure 6 (g)), accompanied by the formation of long torturous segments between the branching points (Figure S18 (c)) and resulted in increased total blood vessel density (Figure S18 (a)). At the same time the elongation of the branching blood vessels in CAMs grafted with TECs and scaffolds was rather inhibited as demonstrated in Figure 6 (c) and 6 (e). The combination of these changes resulted in formation of the areas of denser blood vessels networks between the proper branching points. Figure 6 (f) shows the empirical cumulative distribution functions (ECDFs) visualizing the branch length density distributions in intact CAMs on *ED8* and in each of 4 groups on *ED12* which summarize the data presented in Figures 6 (a-e) and S18 and quantified in Table S6. These results show that compared with the measurements on *ED8*, the CAMs that developed naturally underwent detectable angiogenesis, while the AOSSes did not enhance this. At the same time the TECs supported the development of higher branch length densities in CAMs than the naturally developing embryos and the unseeded scaffolds (Figure 6 (g)). The most pronounced increase in the branch length density was detected in the *Cells* group. Finally, as shown in Figure S18, in TECs-grafted CAMs, the total blood vessel density and segment length per area were increased in comparison to all other groups.

**Figure 6.**
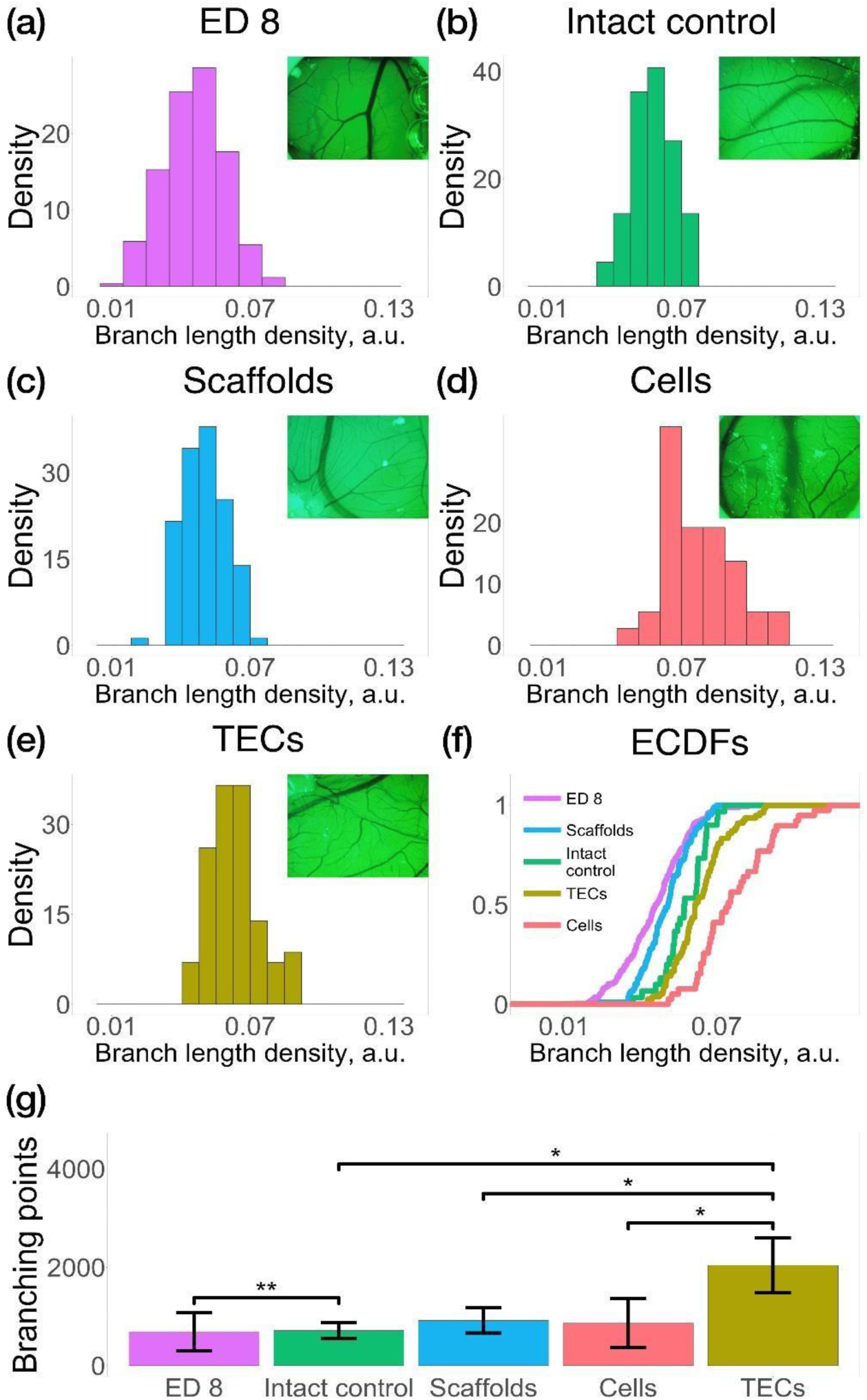
The angiogenic effects of 3D TECs, liver AOSSes (Scaffolds), and suspension of MDA-MB-231 cells (Cells), grafted on a chick embryo CAM compared with natural growth of CAM vasculature (Intact control) in the period between ED8 and ED12 of the embryonic development. Histograms of *branch length density* of (a) CAM vasculature on the day of grafting (ED8), and in (b) Intact control CAM (ED12), (c) Scaffolds, (d) Cells and (e) TECs. The inserts are images of blood vessels located in the vicinity of grafted materials. (f) ECDFs of branch length density in CAMs on ED8 and in each group on ED12. (g) Comparison of the number of branching points (per unit area) in CAM blood vessels in the studied groups. Error bars indicate standard deviation. Stars denote the level of statistical significance by Mann-Whitney test * p<0.05, ** p<0.01.

## DISCUSSION

In this study, we investigated a new model of early metastasis of TNBC to the liver established by seeding acellular solid scaffolds prepared from whole livers of chick embryos by an original non-perfusion iWO-DCL method with TNBC MDA-MB-231 cells. The chick embryo tissue engineering platform for disease modelling is inexpensive, highly reproducible, avoiding the complexities of perfusion-based WO-DCL, and it allows to generate whole-organ scaffolds by a simplified immersion-agitation procedure. The resulting iWO-DCL scaffolds can be cut into smaller fragments (AOSSes), allowing high-throughput assays while preserving unique spatial and chemical organization of liver ECM. Importantly, the model allows maintaining the engineered 3D tumors *in vitro* for long periods (at least 1 month, as demonstrated in this work), which is important for metastatic cancer research. The TECs may also be transferred to *in vivo* settings as tumor xenografts, offering a unique possibility to observe intrinsic mechanisms behind metastatic cancer outside and inside the body. The DCL methodology introduced here maintains the liver-specific ECM histoanatomy with identifiable parenchymal and stromal compartments. It also preserves major molecular components of the ECM such as fibrillar collagen indicated by Van-Gieson’s staining as previously reported for perfusion WO-DCL of the liver ^33, 46^-^48, 50^. According to our pilot experiments ^51, 52^, immersion-agitation decellularization applied to small whole organs or sections of large organs (following the protocol optimization), allows to obtain AOSSes of the quality and biocompatibility comparable with the WO-DCL scaffolds produces by perfusion in bioreactors. Therefore, iWO-DCL liver AOSSes form biologically accurate scaffolds reproducing the hepatic microenvironment more closely in comparison with synthetic scaffolds and culture plastic, both lacking natural signalling moieties and tissue architecture.

We found that *upon first contact with the liver ECM*, TNBC cells preferentially attached to the matrix of the stromal compartments, while their initial adhesion to the ECM of the parenchymal compartments was ∼2.8 times less frequent. Next, *during the first two weeks* of culturing, the cells in the parenchymal compartments mainly remained individual and they dispersed in the spongy-like ECM of former parenchyma, while the stromal compartments were populated mostly by multicellular clusters which lined along denser connective tissue elements. The observed cellular distribution is consistent with the reported finding that the earliest detection of non-vascularized hepatic metastases which presented as cellular clumps in Doppler sonography was possible near the portal triads (i.e., in the stromal compartment) ^53^. It also raises the question about the detectability of single-cell occult metastases in the liver parenchyma. Moreover, during *the first two weeks* of observations, the individual vs. clustered cell adhesion in the compartments was reflected by their subsequent differential invasion patterns.

The features of TNBC cell invasion in the parenchymal compartments can be attributed to *amoeboid migration* ^54^, as the cells mainly preserved their round epithelial morphology and appeared to be separate from each other, permeating through small ECM voids without visible signs of matrix degradation. The stromal compartments were initially invaded by clusters of cancer cells, therefore indicating that *collective cell migration* ^54^, was the early mechanism of the colonization in the stromal ECM. These alternative colonization strategies resulted in different depth of the invasion fronts in the parenchymal and stromal compartments, found to be 134 µm and 70 µm, respectively, implying that individual cell migration in the former hepatic parenchyma resulted in faster speed of the invasion front (∼10 µm/day), while the stromal parts of the liver matrix were invaded more slowly (∼5 µm/day) by TNBC cellular cohorts. This is consistent with the *in vitro* data indicating that individual cancer cells migration is faster than the collective mode due to different adhesion mechanisms and reactions to spatial confinement ^55^. At the same time, the cellular density in the parenchymal compartment by the end of the second week after seeding remained 2.4 times lower than in the stromal zones, indicating, probably, that proliferation and migration states of the cells were, to an extent, mutually exclusive. We conclude that early TNBC colonization of the two liver compartments occurs via different mechanisms and may require different therapeutic approaches (such as cell immobilization in the parenchymal compartment and a cytostatic treatment in the stromal part).

From *Week 3* onwards, after significant remodelling of the stromal ECM causing its loosening and gradual blurring of a structural difference between the liver ECM compartments, the TECs were invaded to the depth of 800 - 1,500 µm from the surface, while the cellular population comprised a mix of cells with epithelioid and mesenchymal-like morphologies. At this stage, a significant increase of the invasion front speed is observed (∼26-50 µm/day) associated with reversible phenotype transitions and matrix remodelling employed by TNBC cells as colonization strategies. This is in agreement with the reported findings of interdependence of the migration mode and morphology of the cells, and transition between the mesenchymal and epithelial phenotypes in the migrating cellular clusters resulting in individual dissemination of epithelial-like cells ^56^.

Our results revealed dramatically different cell population dynamics in the 2D culture of MDA-MB-231 cells and in 3D TECs. Approximately 85% more cells initially attached to the 2D plastic substrates then to AOSSes, and the plastic-cultured cells grew much faster during Week *1* than cells in the 3D TEC. Later, in 2D culture, the growth rates stabilized and even decreased in *Week 4*, while in 3D TECs the cell density was steadily increasing during the *first 3 weeks*. We found that a logistic growth model was applicable to both cultures; such model accurately captures growth of breast carcinoma cells in large clinical datasets ^57^. Logistic growth indicates the presence of resource limitations ^58^. In 2D cultures, this was likely to be due to saturation of the cell attachment capacity of the wells ([∼ 1.9×10^5^] cells, the value reached in Day 2 – 3, Figure 3 (a)). This was not the case for the TECs because of the initially smaller numbers of attached cells, and large surface area of the porous ECM scaffolds. We attribute the decrease of cell growth rate in TECs after *Week 3* to metabolic limitations of 3D cell culture, including hypoxia ^59^. This agrees with our histological observations of secondary necrotic areas forming at the late stages of TECs development.

Our study demonstrates the feasibility of the proposed tissue engineering TNBC model for drug and nanomedicine research. We found that the sensitivity of TNBC cells to the chemotherapeutic drug doxorubicin in free and nanoformulated forms was significantly compromised in 3D TECs, in comparison to 2D cultures (Tables S4, S5), with 9-times increase of EC50 values and the value of over 10 µg/mL for IC50. This is in agreement with earlier reports that additional signalling inputs from the 3D microenvironment and ECM ligands lead to decreased responses to chemotherapeutics ^60^. We found higher therapeutic effect of the nanoformulated Dox in 3D TECs in comparison with free Dox, notable at high doses of doxorubicin (Figures S14) with significant difference at 10 µg/mL (p < 0.03). This result was corroborated by fluorescence imaging, where an uptake and intracellular distribution of free and nanoformulated Dox were found to be comparable in 2D TNBC cultures (Figure 5 (a-h)), while in 3D TECs the AMS-6-Dox nanoparticles killed deeply located cells almost completely, in contrast with the free Dox, which accumulated in the cell nuclei, but did not induce tumor decay (Figure 5 (i-p)). Therefore, this difference is tentatively attributed to different diffusion characteristics and/or hypoxia potentiation of the cytotoxic effect of Dox in deeper layers of TECs.

Finally, we validated our tissue engineering tumor model *in vivo* by an angiogenic assay on chick embryo CAM. Angiogenic switch is the key limiting process for a transition between micro- and macrometastatic states ^2^. We found that grafted TECs induced an enhanced angiogenic response in comparison with natural embryonic CAM vascularization. In contrast to natural embryonic angiogenesis, which occurs within the studied incubation age mainly by *intussusceptive microvascular growth* (i.e. splitting of the existing capillaries along the blood vessel axis) ^61^, the grafted tumor stimulated branching of the host blood vessels, indicating that *sprouting angiogenesis* was taking place. Interestingly, the increase of branching, in contrast to the increase of blood vessel length density, in comparison to controls, was not induced by grafted cell suspensions, indicating the contribution of the ECM-related factors on the tumor-specific angiogenic potential of the TECs. As the sprouting angiogenesis is the key part of the angiogenic switch in the metastatic progression ^2, 62^, this result indicates that the presented model of early TNBC liver metastases behaves similarly to early metastasis.

In conclusion, the new engineered 3D metastatic cancer model investigated here fills the gap between conventional cell cultures and *in vivo* testing in animals, and it provides an affordable and facile platform for drug development and cancer research. This model allowed us to reveal distinct, stage-specific and previously unknown strategies of metastatic colonization of TNBC in the liver. The compartmentalization of the liver ECM was found to be the key determinant of the colonization pattern employed by TNBC cells. Cells cultured in 3D liver-derived matrix demonstrated increased resilience to free and nanoformulated doxorubicin. The biological relevance of the model was confirmed *in vivo* by induction of angiogenic switch in the host tissue by the grafted TECs. As decellularized tissues have negligible cross-species differences and can be used as xenotransplants ^32^, this methodology is highly universal and may be applied to other types of chick embryo organs and other cell lines.

## Supporting information

Manuscript

## ACKNOWLEDGEMENTS

We gratefully acknowledge discussions and critical review of histological results by Professor Anatoly Shekhter (Sechenov University). We are thankful to Ms. Larisa Grivans for her assistance with scientific illustrations. A gift of fertilized chicken eggs from Baiada Poultry PTY Ltd is thankfully appreciated. This work was partially funded by the ARC Centre of Nanoscale Biophotonics CE14010003 and by the Ministry of Education and Science of the Russian Federation (project No. 14.Z50.31.0022). AG is grateful to Macquarie University for providing an IMQRes scholarship.

## CONFLICT OF INTEREST STATEMENT

We confirm that confirming that all experiments were performed in accordance with relevant guidelines and regulations. The study protocol was approved by the animal ethics committee of Macquarie University, Sydney, Australia (ARA 2015/006). The authors declare no competing interests.

The datasets generated and/or analysed during the current study, as well as the custom-built computer codes are available from the corresponding authors upon a reasonable request.

## AUTHORS’ CONTRIBUTIONS

The authors contributed the following to the preparation of this paper:

Concept and design – A.G., A.Z.

Planning and implementation – A.G. A.Z., E.G., Y.Q.

Data collection – A.G., I.K., V.R., Z.K., A. G.-B.

Analysis and data interpretation – A.G., V.R., I.K., Z.K., A.N., L.L., E.G., A.Z., Y.Q.

Writing the manuscript, preparation of the figures and artworks – A.G., V.R., I.K., A.N., L.L.

Editing of the manuscript – E.G., A.Z., A.G., A.N., A.G.-B., Y.Q. Joint co-supervision of the project – A.Z., Y.Q. and E.G.

## METHODS

### Cell culture

MDA-MB-231 (ECACC 92020424) cells were expanded by culture in complete culture medium prepared from Dulbecco’s Modified Eagle’s Medium (DMEM/F12/Ham medium, #D8437, Sigma-Aldrich) supplemented with 10% fetal bovine serum (FBS, #12003C, Sigma Aldrich) and 1% Penicillin-Streptomycin (PS, 10,000 U/mL; #15140122, Gibco) under standard conditions (37 °C, humidified, 5% CO_2_ gas atmosphere) during 7-10 days prior use in further experiments to reach the 4^th^ – 6^th^ passage. Culture medium was changed every two days and the cellular growth was controlled by using a phase-contrast microscope and cell counting. According to the cell counting data, the average population doubling time of MDAMB-231 cells (passages 4 to 6) was approximately 34 h, with the average viability ∼98%. The same culture medium was used for all *in vitro* experiments, unless otherwise specified.

### Chick embryo incubation and organ collection

The study was approved by the animal ethics committee protocol of Macquarie University (ARA 2015/006). Fertilized chicken (Gallus gallus domesticus, the strain of White Leghorn) eggs were delivered from a local hatchery, allowed to rest for 3-4 h at room temperature and then incubated in a standard cradle-type laboratory poultry incubator (R Com MARU Max 190, Autoelex Co., LTD, South Korea) at 37.5 °C, 65-70% humidity with hourly turn over until the embryonic day 18 (ED18), 3 days before natural hatching. This period was sufficient for the development of the organ structure and main physiological systems of a chick, excepting the central nervous system. The pain sensitivity was immature at this stage, and it was permissible to extract an embryo from the egg for research purposes in accordance with the Australian code of animal research. Egg shells were opened by a cut on the blunt ends of eggs, and the embryo with embryonic membranes were extracted using forceps and immediately decapitated. Then feathers were removed from the abdominal wall and thorax, and the liver was carefully extracted through the wide central section.

### Decellularization

The iWO-DCL was performed by our original method ^42^ with some modifications. The extracted chicken embryo livers were washed in sterile PBS and placed in 50-mL Falcone tubes, 5 livers/ per tube filled with 30 mL of 0.1% solution of sodium dodecyl sulphate (SDS) in phosphate buffer saline (PBS), then closed tightly and fixed horizontally on the platform of an orbital shaker. Then the organs underwent shaking at the speed of 90-150 rotations per minute (rpm) with periodic aseptic changes of washing media for a fresh portion every 3 h during first 12 h, and then every 6 h during the next 12 h. Afterwards, the solution was changed daily until the organs became translucent and the liquid media turned colourless and transparent. The total processing time ranged from 14 to 21 days depending on the embryo flock. Next, the processed organs were aseptically placed in sterile containers and washed with 1% of antibiotic-antimycotic solution (A-A) (#A5955, Sigma-Aldrich) in PBS (pH 7.0) for 35 days under shaking (30 – 90 rpm) periodically changing A-A/PBS with fresh portions, until the washing media became transparent, colourless, with no observable tissue components and foam (Figure S2). The processing was performed at room temperature. As-obtained scaffolds were stored in fresh sterile 1% A-A/PBS solution in a fridge (+4 °C) until further use.

### Recellularization of acellular chick embryo liver scaffolds with MDA-MB-231 cells

#### Preparation of scaffolds (AOSSes) for cell seeding

Small fragments (approximately, 3 × 4 mm) of decellularized livers were cut by a scalpel blade and put into 24-well flat bottom tissue culture plates (Costar, Corning, #3524). 1 mL of 0.1% peracetic acid solution (#77240, Sigma-Aldrich) in 4% ethanol was added to every well and placed onto an orbital shaker platform for 2 h at 50 rpm. Then this solution was replaced with sterile PBS (0.4 mL per well) and decellularized liver AOSSes were sterilized by ultraviolet light in a tissue culture hood for 45 min. Next, PBS was removed, and each well was refilled with 1 mL of complete culture medium. Following that, the plates with the AOSSes were placed into a tissue culture incubator and conditioned overnight under humidified atmosphere with 5% CO_2_ at 37°C.

#### Seeding cells on liver AOSS to prepare liver-specific tissue engineering constructs (TECs)

MDA-MB-231 cells (1 × 10^5^ cells in a 30-µL drop of complete culture media) were seeded on as-obtained AOSSes, one TEC per a well of 24-well culture plate. Control scaffolds were left unseeded. Next, the cells were allowed to attach to the substrates for 2 h in a tissue culture incubator, then added with 1 mL of complete culture media per well and cultured for 1-28 days. Medium in these growing cultures was carefully changed twice a week. TECs were sampled for imaging using a fluorescence microscope, performing viability assay and histological analysis on weeks 1, 2, 3 and 4 after seeding.

### Morphological study of in vitro structural evolution of TECs

AOSSes were sampled and investigated after the DCL procedure to examine their structural integrity and sterility and as-developed TECs were also sampled for morphological study after 1, 2, 3 and 4 weeks of incubation with cancer cells. The samples were fixed in 10% neutral buffered formalin, dehydrated in a graded series of alcohols, embedded in paraffin wax and cut into serial sections of 5 µm in thickness by a rotary microtome. After thorough deparaffination, slices were stained with haematoxylin and eosin (H&E), van Gieson’s picrofuchsin, Masson’s trichrome and toluidine blue, following conventional protocols. In addition, unfixed TECs were stained with 4’,6-diamidine-2’-phenylindole dihydrochloride (DAPI) to highlight the cell nuclei. Staining with fluorescein diacetate (FDA) and propidium iodide (PI) was used to label and discriminate live and dead cells, respectively. Stained histological preparations were examined using an upright research microscope Axio Imager Z2 (Zeiss, Germany) equipped with dry-air EC Plan-Neofluar (5×/NA0.16; 10×/NA0.30; 20×/NA0.50 Ph) and oil-immersion α Plan Apochromat (100×/NA1.46 oil) objectives (Zeiss, Germany). Fluorescence microscopy was performed within 30 minutes after collection and staining of the samples with the use of the filter settings for DAPI, and FITC (for FDA) and PI on the same microscope. The images were recorded using a preinstalled microscope digital video camera AxioCam (1388×1040, Zeiss, Germany) in a single-frame and stitching modes and analyzed using Zen 2012 proprietary software.

### Histological morphometry

Image processing techniques were applied to evaluate the H&E stained histological images of the evolving TECs acquired on *Weeks 1, 2, 3*, and *4*. During pre-processing, Gaussian blur with s.d. σ = 0.65 μm was applied to reduce high-frequency components. Using colour deconvolution the images were split in three separate channels containing cells, matrix, and background respectively. Next, images with cells were thresholded and segmentation algorithm was performed where possible to extract single cells boundaries. Cell shape was described by calculating the circularity of a convex hull. The reason for using convex hull approximation of cell boundaries is that it reduces variability of the data and allows more accurate classification.

Image processing was performed using ImageJ and MatLab 2016b.

### MTT assay for evaluation of the population evolution of MDA-MB-231 cells in 2D and 3D TECs

The MDA-MB-231 cell viability was tested in 3D TECs and in matching 2D cell cultures using a modified MTT colorimetric assay. This assay relies on the reduction and conversion of yellow 3-(4,5-dimethylthiazol-2-yl)-2,5-diphenyltetrazolium-bromide (MTT) reagent (#M2128, Sigma-Aldrich) into purple formazan salt, where the optical absorbance of formazan crystals dissolved in dimethyl sulfoxide (DMSO) represents the activity measure of cellular mitochondrial dehydrogenase ^63^. The tested cultures were grown in complete culture medium in a humidified atmosphere under 5% CO_2_ at 37°C.

The following procedure was introduced to ensure appropriate comparison between 2D and 3D TEC cultures. The 5th passage MDA-MB-231 cells were seeded on chick embryo liver AOSSes, as described earlier, while the same amount of the cells (1×10^5^ in a 30-µL drop of complete culture media) was deposited in the middle of 24-well culture plate (Costar) wells to perform high-density seeding. Next, cells in both cultures were allowed to attach to the substrates for 2 h in a tissue culture incubator in a humidified atmosphere under 5% CO_2_ at 37°C, and then filled with 1 mL of complete culture media per well and cultured for 1 day.

After 24 h, the media was removed, and the samples were washed twice with PBS to eliminate unattached cells. Next, 3D TECs were aseptically transferred to new 24-well culture plates to get rid of the cells adhered to the plastic and not to the scaffolds in the original cultures, then filled with fresh complete culture media (1 mL per well) and cultured for 4 weeks. At the same time, 2D cell cultures after washing with PBS, were filled with complete culture media, and cultured for 4 weeks, without splitting, in the same way as 3D TECs. The media was changed twice a week in both types of cultures.

The MTT assays were carried out on days 1, 7, 14, 21 and 28 after seeding (day 1 and *Weeks 1, 2, 3 and 4*, respectively). For each assay, 3 samples of TECs were randomly selected and transferred to a separate 24-well plate for testing, while 3 wells of cells growing in 2D culture were used as the internal control. After double washing with PBS, 500 µL of MTT reagent (0.5 mg/mL in the phenol red free cell culture medium; DMEM/F12; #D6434, SigmaAldrich), was added to each well. Then the samples were incubated at 37 °C in a tissue culture incubator for 1 h to allow precipitation of insoluble formazan crystals. After that, the supernatant was carefully collected and 500 μL of DMSO was added to the wells and left for 10 min in the dark on a rocking platform at room temperature to dissolve purple formazan crystals. Next, four portions of 100 µL of the dissolved MTT product was taken from each well, transferred to separate wells of a clear 96-well culture plate (#3585, Costar, Corning) and used for absorbance measurements. The samples’ absorbance was measured in a spectral band centered at 570 nm by a PHERAstar multiplate reader (BMG Labtech, Germany), with empty wells used as blank controls. Each reading was repeated twice; the results were corrected for the blank controls by MARS Data Analysis software (BMG Labtech, Germany).

The average number of viable cells in 2D cell culture on day 1 (24 h after seeding) was calculated by taking into account the population doubling time in 2D cell culture *in vitro* and the seeded number of cells, and resulted in *N*_1_ = 163×10^3^ cells per well. Since the absorbance is proportional to a number of viable cells, the mean number of cells *N* in each group and the standard deviation *σ*_*N*_ were calculated as follows:

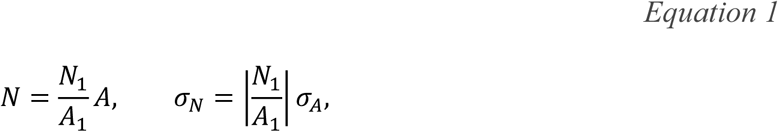

where *A*_*1*_ is the average absorbance in 2D cell culture on day 1, *A* is the average absorbance in each group, and σ_*A*_ is a standard deviation of the absorbance in each group.

The mean increase in cell population per week Δ*N* and the standard deviation *σ*_Δ*N*_ were estimated in both cultures according to the following formulas:

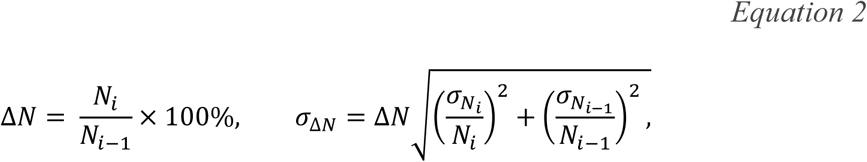

where *N*_*i*_ is the number of cells in week *i* and *N*_*i*−1_ is the measurement in the previous week, *σ*_*Ni*_ and *σ*_*Ni*−1_ are standard deviations of the cell numbers, respectively, in week *i* and in the previous week.

### Modelling cell growth dynamics

The logistic growth model was fitted to the data containing cell numbers at each point in time in 2D and 3D cultures:

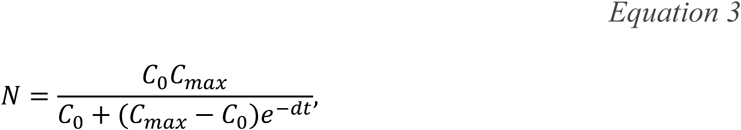

where *N* is a number of cells at time *t, C*_0_ represents the initial cell number, *C*_*max*_ is a maximum number of cells, and *d* is a constant growth rate. The results of the fitting and the estimated values of the model parameters are provided in TablesS2 and S3, respectively. The fit was performed using nonlinear least squares function in R software.

### Cytotoxicity and cellular uptake of doxorubicin (Dox) and mesoporous silica nanoparticles loaded with Dox in 3D TECs and 2D cultures

#### Preparation of mesoporous silica nanoparticles and their loading with Dox

Anionic surfactant-templated mesoporous silica nanoparticles (AMS-6) have been reported by us in Refs ^64, 65^. Nanoparticles were synthesized in-house following a protocol described elsewhere ^65^. Briefly, N-lauroyl-L-alanine was used as surfactant, APES was applied as a costructure directing agent to achieve connected pores in TEOS-sourced silica nanomaterial. The sample was calcinated at ∼550°C using the temperature gradient of 1.5°C/min to remove surfactant. Next, as-synthesized mesoporous silica nanoparticles (AMS-6) were loaded with 20% Dox (Doxorubicin hydrochloride, #D1515, Sigma-Aldrich). Dox diluted in 100% ethanol was added to AMS-6 nanoparticles in a round bottom flask mounted on a rotary evaporator, and ethanol was evaporated at 40°C under vacuum with slow rotation. The collected sample was air dried overnight.

#### Characterization of mesoporous silica nanoparticles

Unloaded AMS-6 and Dox-loaded AMS-6 (AMS-6-Dox) were characterized using transmission electron microscopy (TEM), X-ray diffraction (XRD), thermogravimetric analysis (TGA) and dynamic light scattering (DLS). Nitrogen adsorption/desorption isotherm measurements were carried out to evaluate the effective surface area of the AMS-6 and AMS6-Dox samples. For TEM sample preparation, a small amount of dry AMS-6 or AMS-6-Dox was thoroughly crushed in a mortar and then diluted in ethanol. A drop of the suspension was placed on a copper grid and dried. Next, the grid was placed in a sample holder of a JEOL 3000F TEM (Peabody, USA) and imaged at 300 kV with the resolution of 1.6 Å. Images were obtained using Gatan SC1000 11-megapixel CCD camera (Pleasanton, USA), with a 1024×1024 pixel Gatan image filter (Pleasanton, USA). XRD measurements of 20 mg of the dried nanoparticles’ samples were carried out using XRD instrument Bruker D8 Discover equipped with VÅNTEC-500 detector featuring a 140 mm diameter window (Billerica, USA).

XRD patterns were recorded using Cu Kα anode (λ=0.1542 nm), operating at 40 kV and 30 mA. TGA measurements were performed using 1-mg sample placed in an aluminium crucible (TA Instruments TGA2050, New Castle, USA) and heated from 25 to 850°C at 10°C/min under air flow at 10 mL/min. Nitrogen adsorption/desorption isotherms were acquired at a temperature of −196°C using liquid nitrogen with a TriStar II by Micromeritics® instrument (Norcross, USA), following the AMS-6 and AMS-6-Dox samples degassing under vacuum using VacPrep(tm) 061 by Micromeritics® instrument (Norcross, USA) for ∼10 h at 120°C. The surface area was calculated using the Brunauer–Emmett– Teller (BET) equation ^66^.

Hydrodynamic diameters and the zeta-potentials of colloidal AMS-6 and AMS-6-Dox were measured in PBS and complete culture media (1 mg/mL) by Zetasizer Nano ZS (Malvern, UK) in three runs followed by averaging.

#### MTT viability assay of 2D and 3D TEC cell cultures treated with free and nanoformulated Dox

The TECs were cultured for 3 weeks as described above. For control 2D *in vitro* culture MDA-MB-231 cells were seeded in 96-well plates (Costar, Corning, #3599) at the density of 2×10^4^ cells per well and incubated in complete culture medium for 24 h before the test. Then culture medium was removed from all the cultures, TECs were aseptically transferred to new 24-well culture plates and all the cultures were washed 3 times with PBS. Free Dox of the concentrations ranging from 0.1 to 10 µg/mL, AMS-6 (50 µg/mL) and AMS-6-Dox of the concentrations ranging from 0.5 to 50 µg/mL (Dox-equivalent, 0.1 – 10 µg/mL) were diluted in complete culture media and sonicated immediately before the test. Each concentration of each type of the tested compounds was applied in a total volume of 100 µL to 8 parallel wells of 96-well plates for challenging of 2D culture of MDA-MB-231 cells. At the same time, 2 parallel TECs growing in 24-well plates were used for testing of each concentration of each tested compound, and the added volume of the dispersions was 400 µL per a well. The 2D and 3D cultures treated with complete culture medium were used as a control. The exposure time was 36 h. Then MTT tests were performed, as described above (see 2.4.2) with minor changes. In particular, after removal of culture media and washing with PBS, the TECs and cells were incubated in MTT solution in phenol red free culture media (0.5mg/mL) for a longer period of 3 h. Then the supernatant was removed and 100 µl or 400 µL of DMSO was added to the wells of 96-well plates (2D cultures) and 24-well plates (3D TECs), respectively. The solution of formazan in DMSO from the TECs was transferred to the new 96-well plate (100 µL per a well; 3 samples per a TEC), and the absorbance of the tested cultures was measured. The mean percentage of dead cells *E* and the standard deviation *σ*_*E*_ were recalculated based on the absorbance of the controls:

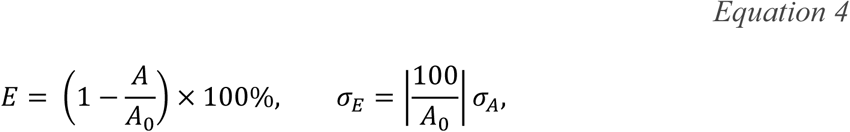

where *A* is the average absorbance in each group, A_*0*_ is the average absorbance of the corresponding control, σ_*A*_ is the standard deviation of the absorbance in each group.

#### Modelling the pharmacodynamics of Dox

The experimental measurements of the percentage of dead cells in 2D and 3D cultures after administration of free or nanoformulated Dox were described by the following sigmoidal equation:

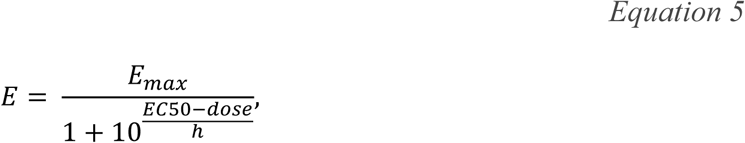

where *E*_*max*_ is the maximum effect (%), *EC50* is the half maximal effective concentration (µg/mL), *dose* is the concentration of administered Dox (µg/mL), and *h* is the Hill coefficient. The half maximal inhibitory concentration (IC50), i.e. the drug concentration needed to obtain 50% of cell death, was calculated in each group using the line of the best fit. The parameters of the model are provided in Tables S5 and S6, respectively, for free and nanoformulated Dox.

The fit was performed using a weighted unconstrained nonlinear curve fit (Matlab r2016b) with the inverse error as the weight, and the dose values were first logarithmically transformed. The 0% cell death (100% viability) at 0 µg/mL in controls was given a higher weight as the data was normalized to these values and the model is expected to approach this intercept closely.

#### Confocal microscopy study of the uptake of free and nanoformulated Dox in 3D TEC and 2D cultures

As produced TECs were cultured for 4 weeks. Control 2D *in vitro* cultures of MDA-MB-231 cells were seeded onto sterile coverslips placed into wells of a 24-well plate (Costar) at the density of 5×10^4^ cells/well and incubated under standard conditions in 1 mL of complete culture media during 24 h prior to the observation. Dox solution (10 µg/mL), AMS-6 nanoparticles (50 µg/mL), and AMS-6-Dox nanoparticles (50 µg/mL; Dox-equivalent, 10 µg/mL) dispersions were prepared as described above. Next, after removal of the culture media and triple washing with PBS the tested compounds were added to the wells with 2D and 3D cultures in a total single volume of 0.5 mL per well. The wells added with complete culture media without Dox or nanoparticles were used as controls. The prepared 2D and 3D cultures were incubated for 24 h in tissue culture incubator at 37 °C an 5% CO_2_, and, next, following thorough rinsing with PBS 3 times to remove free Dox and nanoparticles, they fixed with 10% neutral buffered formalin at room temperature.

After 24-h fixation process and another washing with PBS, the fixed samples were stained with DAPI solution in PBS (#D9542, Sigma-Aldrich) for 20 min at 37 °C. Next, the staining solution was removed, the coverslips were washed twice with PBS to eliminate unbound DAPI.

Finally, the samples were mounted on glass slides with Dako anti-fade mounting media and sealed with nail polish.

Dox and AMS-6-Dox cellular uptake was imaged by an inverted Zeiss LSM 880 laserscanning confocal microscope (Zeiss, Germany), using a Plan-Apochromat 10×/0.45 N.A. M27 and Plan-Apochromat 40×/1.3 N.A. oil DIC UV-IR M27 objectives. Dox fluorescence was observed using 488 nm excitation and emission 535-673 nm; DAPI fluorescence was observed using 405-nm excitation and emission 411-528 nm.

### Evaluation of angiogenic potential of TECs and AOSSes in vivo

#### Grafting and imaging procedures

The detailed description of the used procedures of angiogenic assay on chick embryo CAM^67^ can be found in SI and Figures S16, S17. Briefly, TECs, AOSSes or cell suspensions were grafted on CAM separately, one sample of each material type per egg, 5 replicates per a group. Before grafting TECs were cultured *in vitro* for 12 day as described above, reaching the stage when they contained approximately 2×10^5^ viable cells (according to MTT assay data). The liver AOSSes were kept in complete culture media for 24 h before grafting on CAM. The MDA-MB-231 cell suspensions containing 3.3×10^6^ cells per mL of complete culture media were prepared by trypsinization of the *in vitro* monolayer cultures of 5^th^ – 6^th^ passages and 2×10^5^ cells in 60 µL of complete culture media were grafted on CAMs within a sterile rubber ring. The angiogenic effect induced by TECs, liver AOSSes and suspensions of MDA-MB-231 cells in PBS grafted on CAM was evaluated by stereomicroscopy imaging performed on the day of grafting (embryonic day 8, *ED8*) and on *ED12*, in comparison to natural growth of blood vessels of CAM occurring during the same period of chick embryo development. The digital images were analyzed as described in the “Angiogenesis quantification” subsection below.

#### Angiogenesis quantification

The images of CAM taken using 2×/0.5 N.A. objective were processed using the following methodologies to evaluate the dynamics of the vascular length density and branching of blood vessels in CAMs. For *Control* and *Cells* groups, 10 regions of interest (ROIs) per egg were chosen manually. CAMs grafted with 3D engineered tumors (*TECs*) and liver AOSS (*Scaffolds*) were imaged from the 4 corners around the graft, and then 3-4 ROIs were chosen from every corner to obtain approximately 10 ROIs per egg. Since the vessels have higher contrast in the green channel, the RGB images were split to obtain the green component. Histogram stretching of grey level intensities was performed to enhance the image contrast. Then, a ridge detection algorithm [54] involving convolution with the derivatives of a Gaussian smoothing kernel was used to capture the blood vessels by finding local minima, resulting in a skeleton of a vascular pattern. For each ROI, the total branch length of the obtained skeleton was divided by the area of the ROI and then the results were averaged to get the mean vascular density per egg. Branching points were calculated using the Angiogenesis Analyzer plugin for ImageJ [55] (see SI for the details). Image processing was performed using ImageJ and MatLab 2016b.

### Statistical analysis

The data were expressed as means ± standard deviations (SD), and the 95% confidence intervals (CI95%) for the means were calculated. Due to the non-Gaussian nature of the data the two-sided Mann-Whitney U test was used to evaluate the significance of intergroup differences between the means. The two-sampled Kolmogorov-Smirnov test was used to compare any two observed distributions of branch length density. Statistical significance was reported as follows: *p < 0.05, **p < 0.01, ***p < 0.001 or the exact p-value was provided where possible. All statistical analyses were performed using R Statistical Software.

***

See Supplementary Material for additional methods.

