## Supplementary material for "Tissue engineered model of hepatic breast cancer micrometastasis shows host-dependent colonization patterns and drug responses": Manuscript

#### NATIVE STRUCTURE OF A CHICK EMBRYO LIVER AND THE EFFECT OF DECELLULARIZATION

Chick embryo livers investigated in this work on ED18 had a typical lobular microstructure with vascular and biliary elements which appeared less distinctive than in normal human liver tissue. The interlobular connective tissue sheaths were discernible, after special staining (Figure S1 (a, b)). Stromal elements of the liver tissue contained slightly more fibrous collagen than the hepatic parenchyma (Figure S1 (b, c)). The tissue was completely orthochromatic when stained by toluidine blue, indicating low concentration of acid glycosaminoglycans at this development stage (Figure S1 (d)). The liver was coated with a connective tissue capsule, where subcapsular vascular elements were well differentiated and enlarged and surrounded by loose fibrous connective tissue (Figure S1 (e)). The capsular connective tissue was bright fuchsinofilic when stained by Van-Gieson’s method, indicating high concentration of mature (fibrous, crosslinked) collagen (Figure S1 (f)).

The iWO-DCL procedure preserved the shape of the liver, but its volume decreased significantly and the decellularized organ became almost translucent (Figure S2). The original cells and cellular debris in the DCL liver tissue were no longer observable by histological methods (Figure S3). The hepatic parenchyma transformed into a loose, fine mesh-like matrix formed mainly by the residuals of perisinusoidal ECM of Disse’s spaces (Figures S3). The former septal connective tissue and portal elements (portal arteries and veins) and former central veins were identified in the DCL organ by their denser ECM and morphology. Two compartments of the liver ECM could be distinguished, further referred to as the “parenchymal” compartment comprising a mesh-like matrix of the former parenchyma; and the “stromal” compartment comprising the DCL portal vasculature, interlobular septal stroma and acellular walls of former blood vessels, including central veins and portal arteries and veins (Figure S3). The effect of iWO-DCL on the separate components of the liver tissue is schematically presented in Figure S4. The location of the Disse’s space and the composition of the ECM of the Disse’s space, according to the literature data [1-4], are shown in Figure S5.

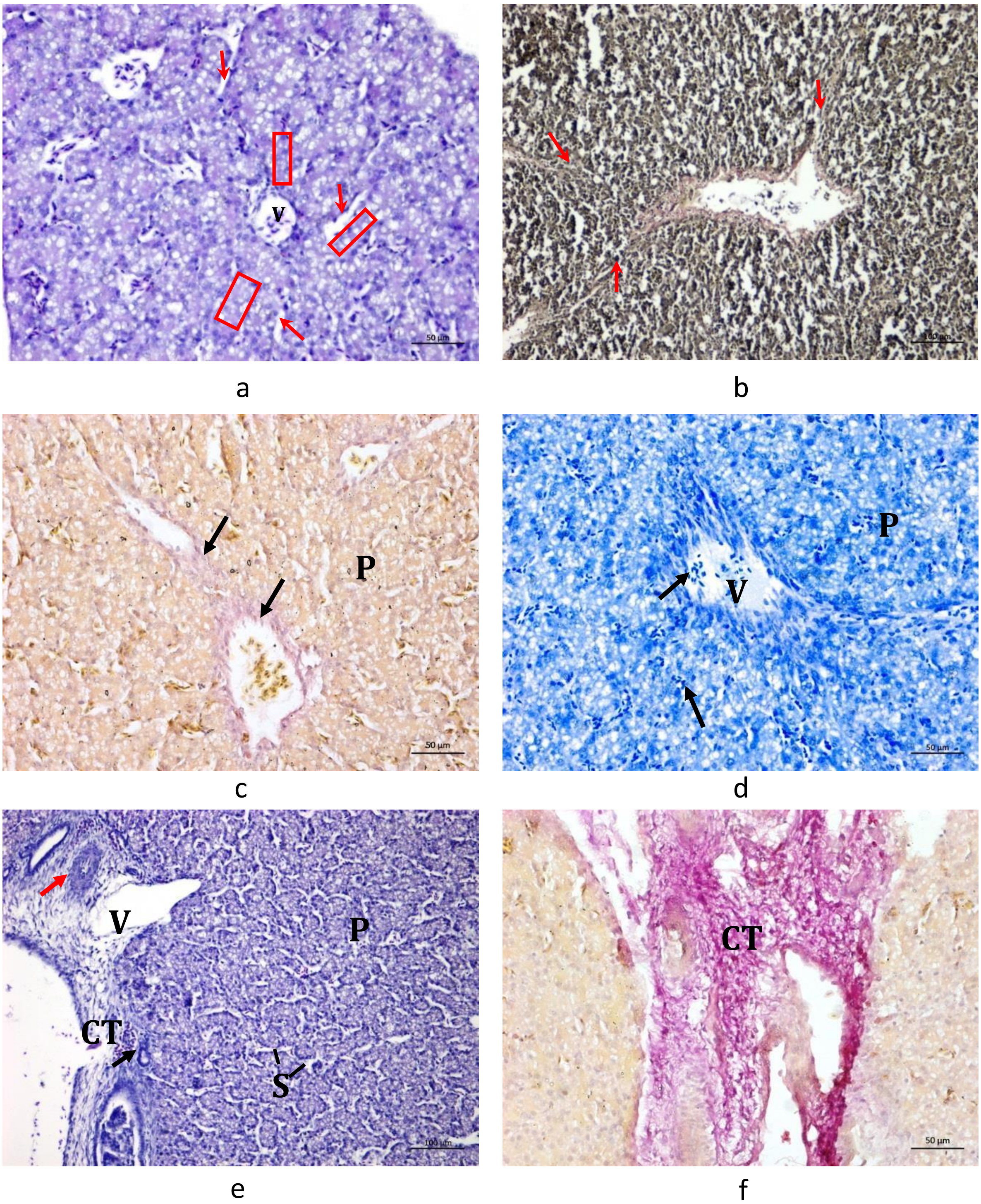


**Figure S1. Details of the histological structure of intact chick embryo liver extracted on ED18.** (a) The cords of hepatocytes (red box) are separated by narrow voids of sinusoids (red arrows) which have clear visible discontinuous endothelial linings, and arranged around the central vein. (b) Lobular structure of hepatic tissue is discernible. Note pink staining of collagen in a large vein and interlobular connective tissue septae (red arrows). (c) Distribution of mature fibrous collagen in parenchymal and stromal elements of the liver tissue. Note weak red staining (fuchsinofilia) of vascular walls (arrows). (d) Orthochromasia of the liver tissue. Note sinusoids containing dark stained nucleated ellipsoid erythrocytes (arrows), which are transported towards the vein (V) through the parenchyma (P). (e) Subcapsular area of the liver. Note connective tissue (CT) of the liver capsule, the veins (V) with wide open lumens of irregular shape and thin walls, an arteria (red arrow) with thicker walls and smaller lumen diameter, and the portal bile ducts with thin walls containing only one row of cubic epithelial internal lining (black arrows). Hepatic parenchyma (P) is formed by cords of hepatocytes separated by sinusoids (S with indicating lines). (f) Connective tissue (CT) of the liver capsule is bright fuchsinofilic. Staining: H&E (a, e); elastica – Van-Gieson's (b); Van-Gieson's (c, f); toluidine blue (d). Scale bars: 100 µm (b, e); 50 µm (a, c, d, f).

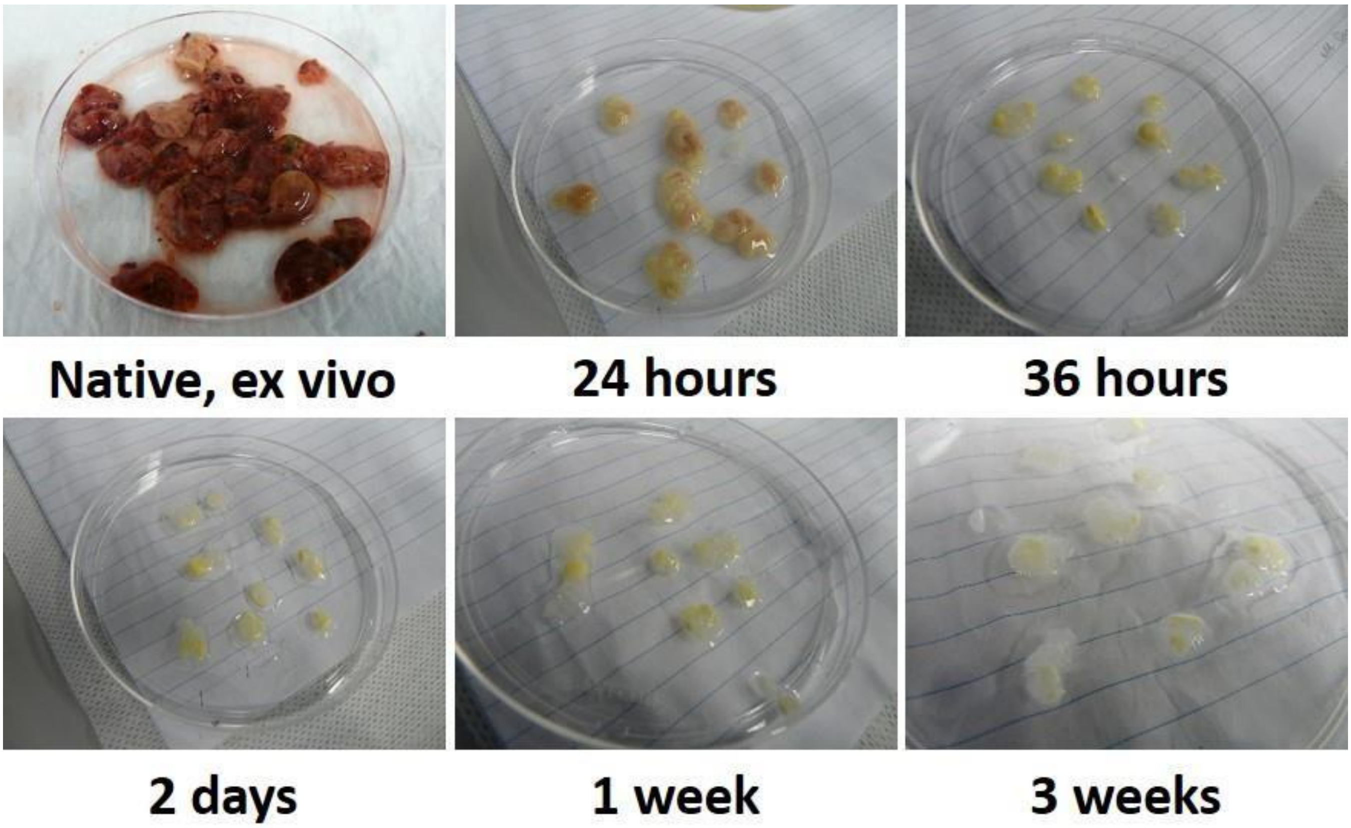


**Figure S2. Effect of iWO-DCL on native chick embryo liver.** Native chick embryo livers collected on ED18 after initial washing in PBS and the macroscopic changes of them during the labelled periods of DCL. Note discoloration and 30% decrease of the organ's volume, comparing to the native state after 24 h of DCL; whitening and decrease of organ's volume to 20% of original after 36 h of DCL; additional down-sizing and emergence of translucent parts after 2 days of DCL; followed by progression of these changes up to the end of the 1st week of DCL. Finally, decellularized livers (3 weeks of DCL) have milky-white translucent appearance and slightly increased volume because of swelling of the loosened tissue.

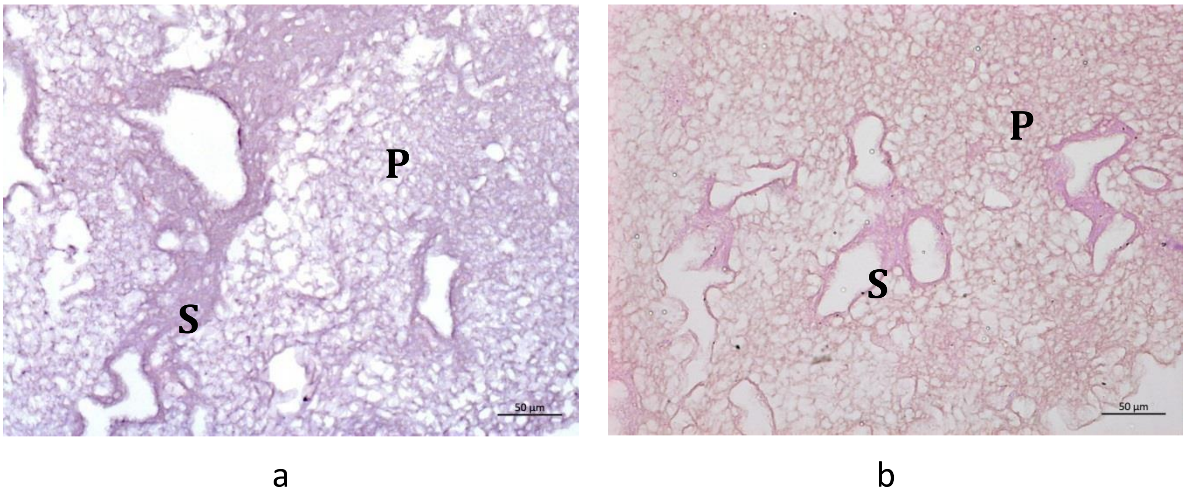


**Figure S3. Effect of iWO-DCL on histological structure of chick embryo liver.** (a, b) Complete removal of cells and observation of compartmental structure of the liver ECM. Note parenchymal (P) and stromal (S) compartments of the chick embryo liver ECM. The ECM of parenchymal compartment is loose and pale stained, with randomly oriented matrix elements of the former Disse's space; the ECM of the stromal compartment is denser, with aligned fuchsinofilic (b) fibrous collagen fibers. The external edges of the scaffolds reveal mild densification of the matrix resulting from immersion-agitation procedure, while there is no distinctive pink fuchsinofilia of these regions in Van-Gieson's stained samples. Staining: H&E (a) and Van-Gieson's (b). Scale bars: 50 µm.

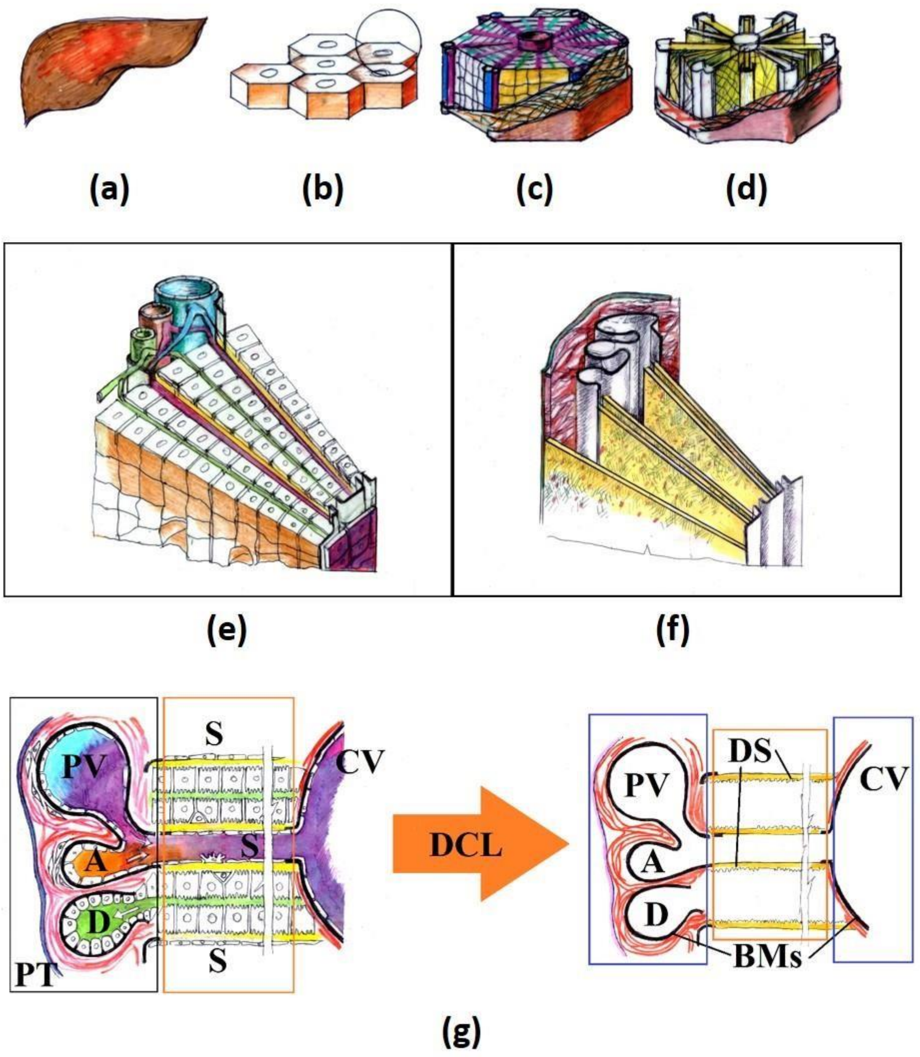


**Figure S4. Schematics of the liver structure and the effect of DCL on its constituents.** The liver (a) is formed by lobules (b). Each lobule (c) has a central vein in the middle and radially arranged cords of hepatocytes separated by sinusoids (radial violet lines) and bile canaliculi (radial green lines). The border of each lobule consists of loose connective tissue sheaths (red coating), which contains portal triads and interlobular vasculature branches in the corners of each lobule's "hexagon". As a result of DCL (labelled as DCL lobule, (d)) all the cellular elements are removed. (e) Detailed view of the sector of a single lobule. Each cord of hepatocytes contains two rows of the polarized cells. One side of the cell row contacts with a sinusoid blood vessel (purple) through the space of Disse (yellow), while the other side of the cell row contacts a bile canaliculus (green) directly. Cells of the sinusoids and other cells of parenchyma, excepting hepatocytes, are not shown for simplicity. Blood from the portal vein (large blue blood vessel) and hepatic artery (smaller red blood vessel) is mixed in the capillaries and transported through the liver parenchyma towards central vein in the middle of lobule. Bile it transported to the portal bile duct in opposite direction. Portal vein, artery and bile duct make a portal triad. (f) 3D view of the effect of DCL on the sector of a single lobule, shown at (e). Note absence of cellular elements and preservation of collagenous stroma of interlobular septae and connective tissue sheaths of large blood vessels (red), basement membranes of large blood vessels and portal bile ducts (grey) and the ECM of Disse's space (yellow). (g) 2D detailed schematic view of a segment of a lobule before (left) and after (right) DCL. Note a portal triad (PT), containing portal vein (PV), hepatic artery (A) and interlobular bile duct (D); sinusoids (S) with discontinuous endothelial lining separated from hepatocytes by Disse's space (yellow). Narrow bile canaliculi (green) are visible between the rows of hepatocytes inside the cords. Bile canaliculi do not have own cell linings. Central vein (CV) has the entire endothelial lining, basement membrane (BM) and collagenous sheath (red lines) around it. The structure of portal blood vessels is similar to that of the CV, but the layer of smooth muscle cells is thicker (it exists only before DCL). Red colour indicates collagen fibres. Black lines indicate basement membranes (BM). "Parenchymal" compartment is shown by orange boxes, and the "stromal" one is indicated by blue boxes.

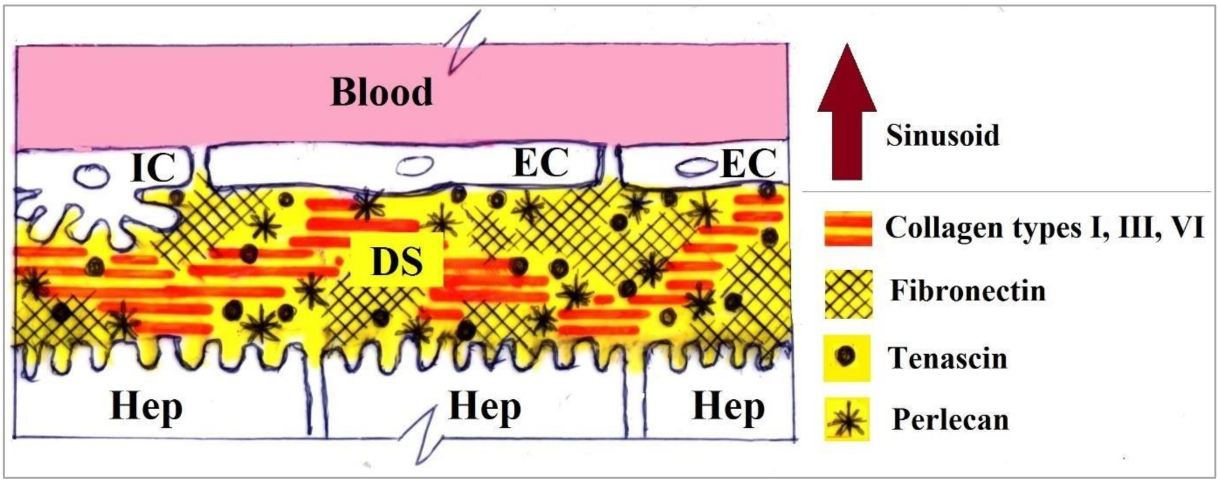


Figure S5. Schematic view of the relationship between sinusoids, Disse's space and hepatocytes (the parenchymal compartment of the liver). Note the wall of a sinusoid formed by discontinuous layer of endothelial cells (EC) and absence of basement membrane underneath the EC. The Disse's space (DS, yellow) separates the endothelial cells of sinusoids from hepatocytes (Hep) and stellate cells of Ito (IC). The concentration of fibronectin increases along the sinusoids towards the central veins. Tenascin is relatively rare component. Perlecan is predominantly associated with basement membranes. Laminin is almost absent along the sinusoids in Disse's space, excepting the area near the portal triads. The scheme is based on References [1-
4].

#### Imaging of TNBC TECs and digital analysis of histopathological data

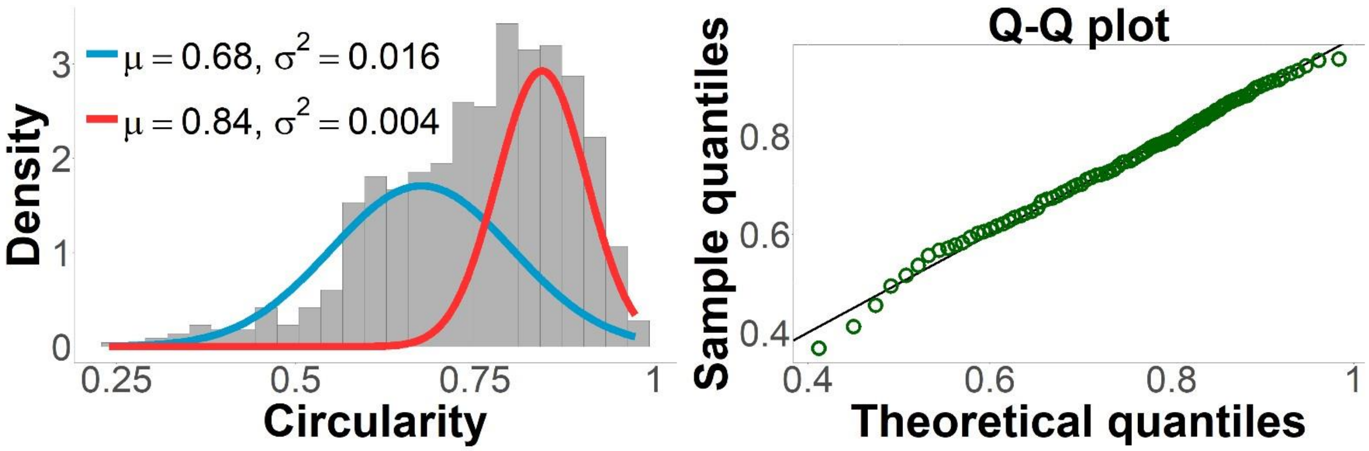


Figure S6. Shapes of individual cells during the first week of culture in the liver ECM microenvironment. Shapes of individual cells were evaluated from histology images obtained in Week 1 by calculating their circularity (μ) calculated as a ratio of the area to the squared perimeter of the cell). Left plot: a histogram of circularity values of the total number of 706 cells. The data is best described by two normal distributions (blue and red curves represent probability density functions) and reveals formation of two subpopulations of cancer cells with different circularities such as elongated oval-shaped "mesenchymal-like" and nearly circular-shaped "epithelioid" cells with circularity values ~0.68 and ~0.84, respectively. Right plot: comparison between the quantiles of the theoretical bimodal normal distribution with the quantiles of the observed data. The data points are positioned close to y = x, indicating that the distribution of the sample data is similar to the normal distribution.

**Table S1. Cell density in different compartments of the liver ECM measured as a ratio of cells to matrix area: results of image analysis of histological sections. 10 images were analysed for each timepoint/compartment.**

| Sampling time point | Compartment | | |
|---|---|---|---|
| | Parenchymal | Stromal | Mixed |
| **Week 1** | 0.051 (±0.069) | 0.163 (±0.152) | - |
| **Week 2** | 0.068 (±0.065) | 0.173 (±0.094) | 0.055 (±0.034) |
| **Week 3** | 0.100 (±0.030) | 0.377 (±0.210) | 0.106 (±0.078) |
| **Week 4** | 0.018 (±0.014) | 0.012 (±NA) | 0.071 (±0.056) |

### CELL GROWTH DYNAMICS IN 3D TECS AND 2D CULTURES OF MDA-MB-231 CELLS

**Table S2. The results of MTT test of viability of MDA-MB-231 cells in 2D and 3D in vitro cultures.**

| Time in culture | Number of cells, $10^6$ | | | |
|---|---|---|---|---|
| | 2D | | 3D | |
| | Mean cell number | 95% confidence interval | Mean cell number | 95% confidence interval |
| **Day 1** | 0.163 | 0.141; 0.185 | 0.024 | 0.022; 0.026 |
| **Day 7** | 1.966 | 1.917; 2.015 | 0.122 | 0.075; 0.167 |
| **Day 14** | 2.491 | 2.357; 2.625 | 0.303 | 0.269; 0.337 |
| **Day 21** | 2.917 | 2.800; 3.034 | 0.454 | 0.389; 0.519 |
| **Day 28** | 2.361 | 2.212; 2.510 | 0.418 | 0.333; 0.503 |

**Table S3. Estimated values of the logistic growth model parameters with standard errors.**

| Parameters | 2D | | 3D | |
|---|---|---|---|---|
| | Estimated value | Standard error | Estimated value | Standard error |
| **$C_0$, cells, $10^6$** | 0.093 | 0.191 | 0.020 | 0.013 |
| **$C_{max}$, cells, $10^6$** | 2.594 | 0.169 | 0.445 | 0.028 |
| **d** | 0.63 | 0.32 | 0.28 | 0.07 |

### CHARACTERIZATION OF NANOPARTICLES

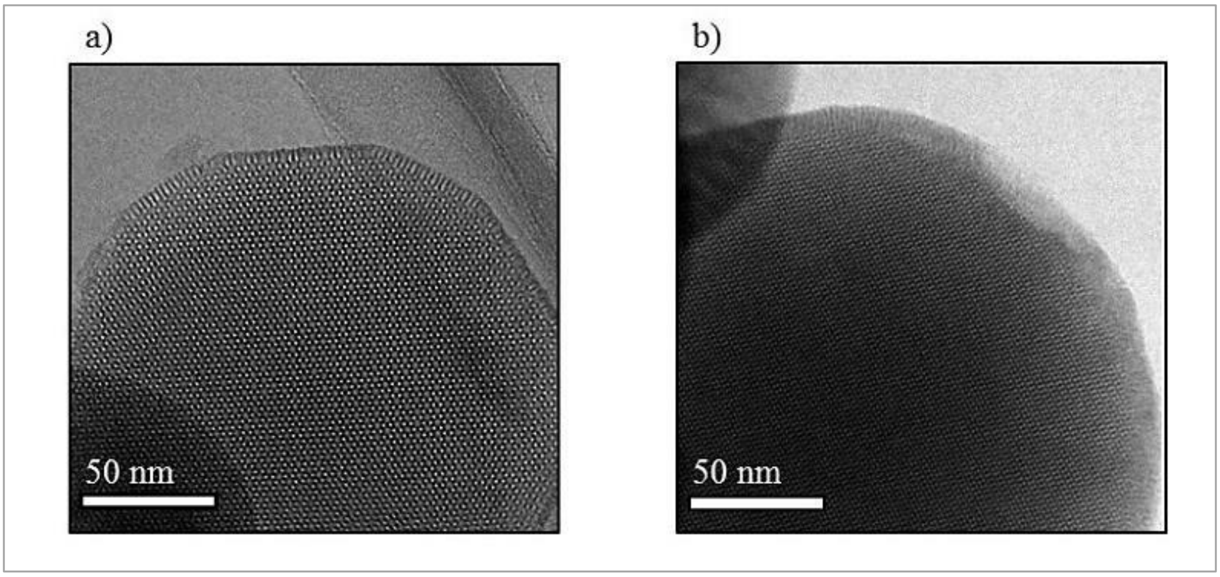


**Figure S7. (a) and (b) Typical TEM images of calcined AMS-6 mesoporous material showing a high degree of mesoporous order.**

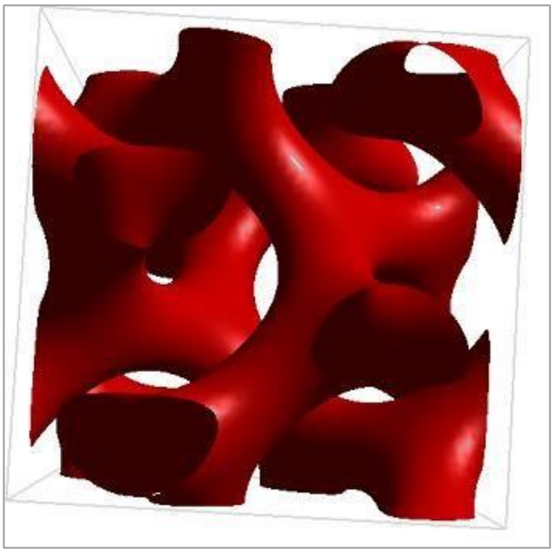

**Figure S8. Schematics of the of AMS-6 structure.**

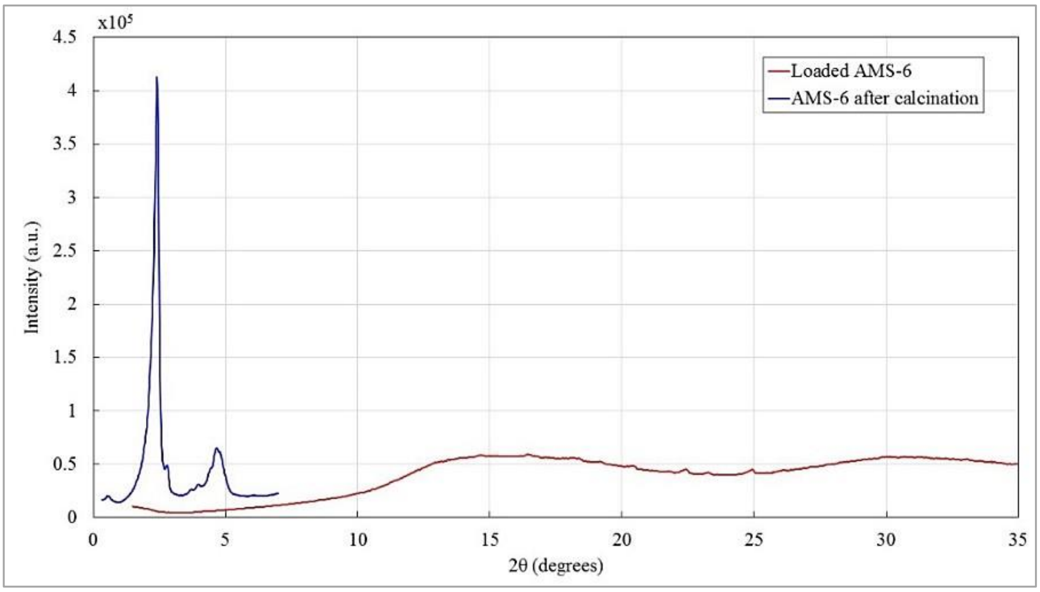


**Figure S9. X-ray diffraction (XRD) pattern of pure calcined AMS-6 mesoporous nanoparticles (blue) and the loaded AMS-6-Dox nanoparticles (red).** The measurements of the pure calcined AMS-6 NPs reveal a highly ordered mesostructure with peaks of scatter in-between the pores of the nanoparticles. The typical XRD pattern of the pure sample display peaks at low angles of 2.4 and 4.8° with an intensity of $4.12 \cdot 10^5$ and $0.6 \cdot 10^5$ a.u., respectively. The loaded AMS-6-Dox sample curve shows small peaks at high angles of approximately 15 and 30° with intensities below $0.75 \cdot 10^5$ a.u., which are most likely the scattering patterns of crystallized Dox outside of the pores. The pores of the AMS-6-Dox still seem to be filled with DOX since the pattern does not show any peaks at low angles which would indicate scatterings between the pore walls of the nanoparticles.

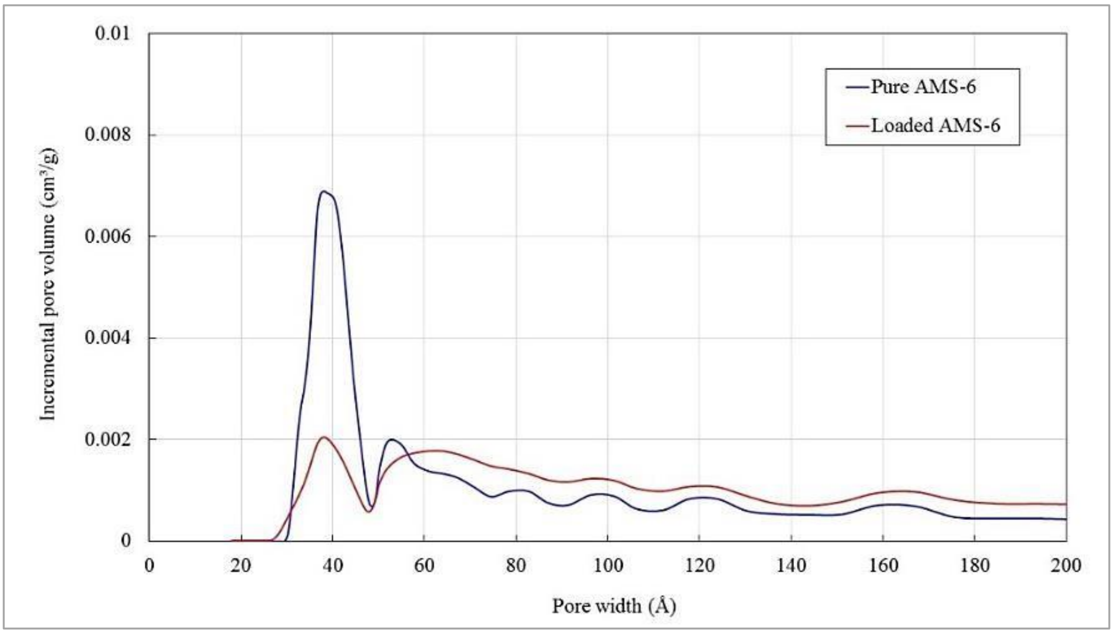


**Figure S10. DFT pore size analysis. The pore width of pure AMS-6 and loaded AMS-6-Dox nanoparticles is in a range of 30 to 50Å.**

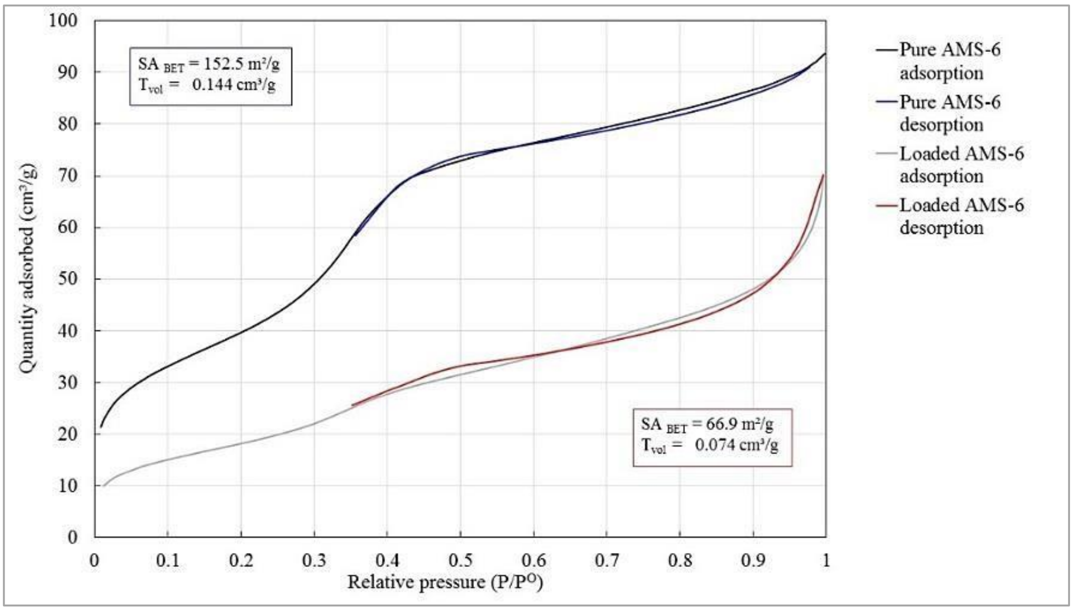


**Figure S11. Nitrogen adsorption/desorption isotherm curves of pure AMS-6 and loaded AMS-6-Dox nanoparticles.** A very narrow hysteresis loop where the plotted adsorption and desorption curves are close together results from the fact that the pore width of 3.81 nm is very small.

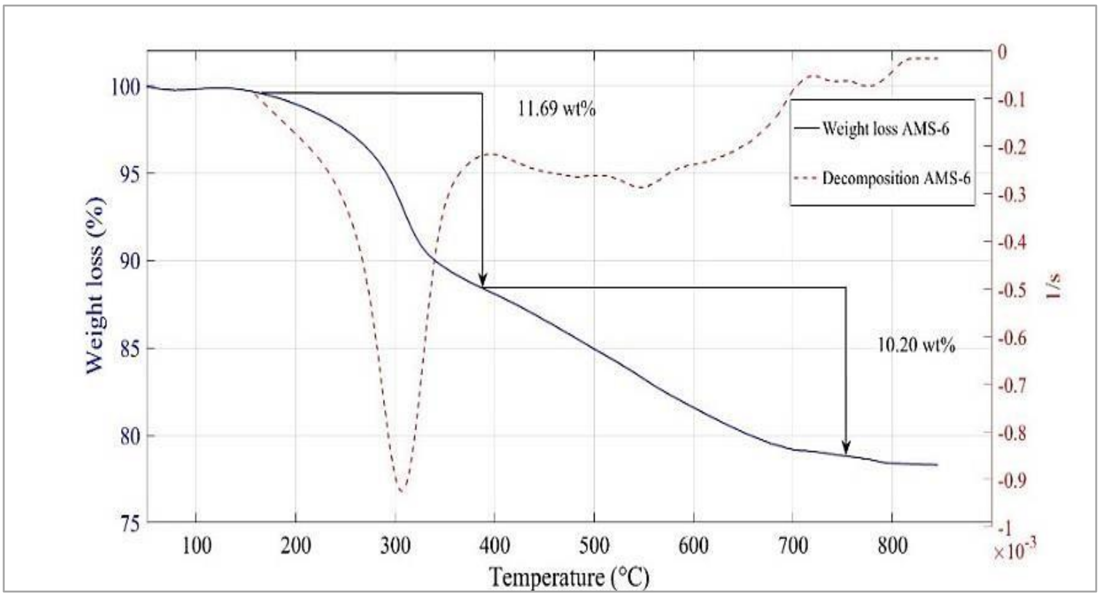


**Figure S12. Thermogravimetric analysis of the unloaded AMS-6 nanoparticles sample.** The thick blue line corresponds to the weight loss versus the temperature, and the dashed red line represents the first derivative from the thermogravimetric analysis (DTG) curve. The latter shows characteristic decomposition peaks, e.g. at 380°C. The measurement was performed on the pure AMS-6 to confirm the absence of any organic compounds that could block the pores, which were supposed to contain the loaded drug later. The peak of DTG at approximately 310 °C corresponds to the loss (decomposition) of propyl amine functionalized groups. The analysis also revealed the incorporation of approximately 11.69 wt% of covalently bound propyl amine groups in the pores of the NPs. However, with 11.69 wt% the presence of amine groups was slightly elevated. A higher value would indicate a high amount of remaining amino groups, which would mean a blockage of the NPs pores. The remaining weight loss above 400°C was due to evaporating water (also known as extra framework water) caused by condensation of silanol groups.

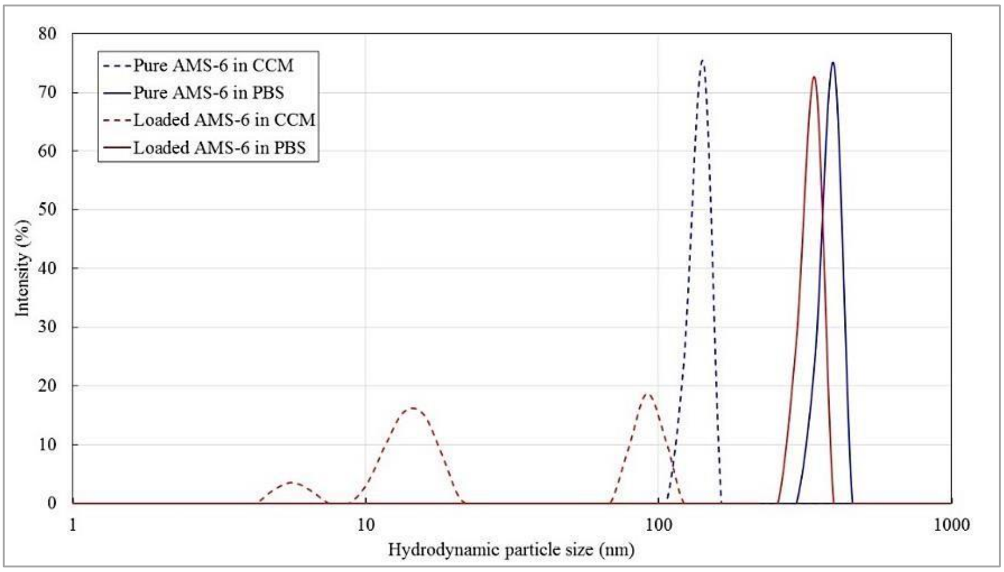


**Figure S13. Hydrodynamic size distribution of pure and loaded AMS-6 NPs in complete culture medium (CCM) and in PBS.** Note the variation of the particles' size distribution depending on the used dilution medium.

### EFFECT OF FREE AND NANOFORMULATED DOXORUBICIN ON MDA-MB-231 CELLS CULTURE IN VITRO IN CONVENTIONAL 2D CELL CULTURE AND IN 3D LIVER ECM-SPECIFIC TECS

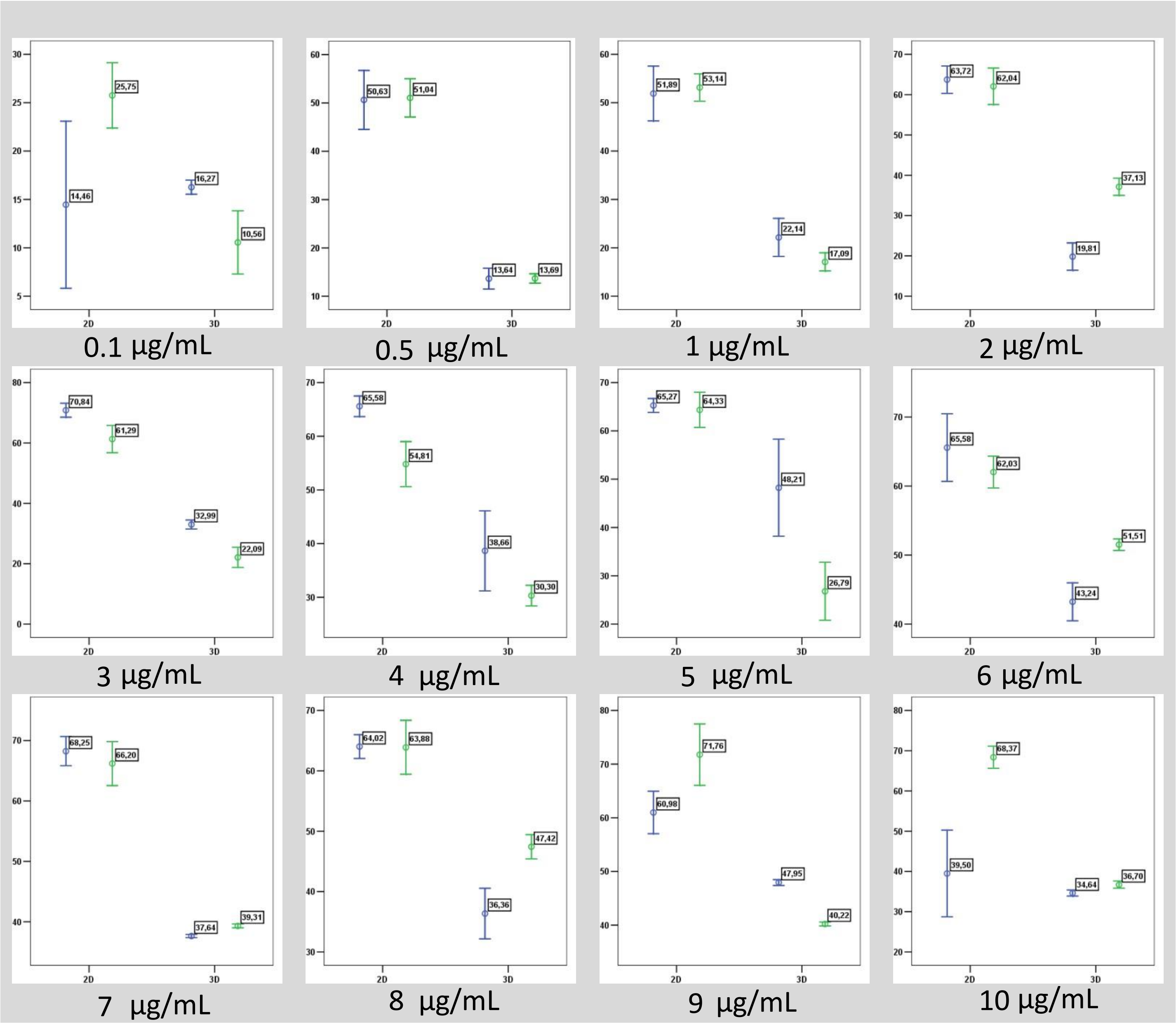


**Figure S14. Intergroup comparisons of the effect of free and nanoformulated Dox in 2D culture of MDA-MB-231 cells and 3D TECs.** Summary of the data obtained by MTT assay in 2D and 3D in vitro cultures of MDA-MB-231 cells exposed to free Dox and AMS-6-Dox nanoparticles during 36 h. Blue and green dots represent the mean ratio of dead cells after application of free Dox and AMS-6-Dox, respectively. Mean ratio of dead cells, % is plotted on Y axis. Experimental conditions, 2D (left) and 3D (right), respectively, are shown by the X axis. Numbers under the graphs indicate the nominal concentration of applied Dox. Error bars show the CI95% for mean percent of dead cells and colour coded in the same manner as the dots standing for the experimental data points. Data labels on the graphs also indicate the mean % of dead cells in the group.

**Table S4. The results of fitting a dose-response model to the measurements of cell viability. Estimated values of dose response model parameters for free Dox in 2D and 3D.**

| Parameters | Free Dox | | | |
|---|---|---|---|---|
| | 2D | | 3D | |
| | Estimated value | Standard error | Estimated value | Standard error |
| **Maximum effect, %** | 64.85 | 2.83 | 34.34 | 2.71 |
| **EC50, µg/mL** | 0.27 | 0.09 | 2.40 | 0.41 |
| **Hill coefficient** | 3.20 | 1.27 | 0.68 | 0.39 |
| **IC50, µg/mL** | 0.43 | 0.12 | > 10 | - |

**Table S5. The results of fitting the dose-response model to the measurements of cell viability. Estimated values of dose response model parameters for AMS-6-Dox in 2D and 3D.**

| Parameters | AMS-6-Dox | | | |
|---|---|---|---|---|
| | 2D | | 3D | |
| | Estimated value | Standard error | Estimated value | Standard error |
| **Maximum effect, %** | 63.06 | 1.97 | 40.53 | 5.47 |
| **EC50, µg/mL** | 0.26 | 0.06 | 2.38 | 0.86 |
| **Hill coefficient** | 2.81 | 0.80 | 0.34 | 0.18 |
| **IC50, µg/mL** | 0.46 | 0.10 | >10 | - |

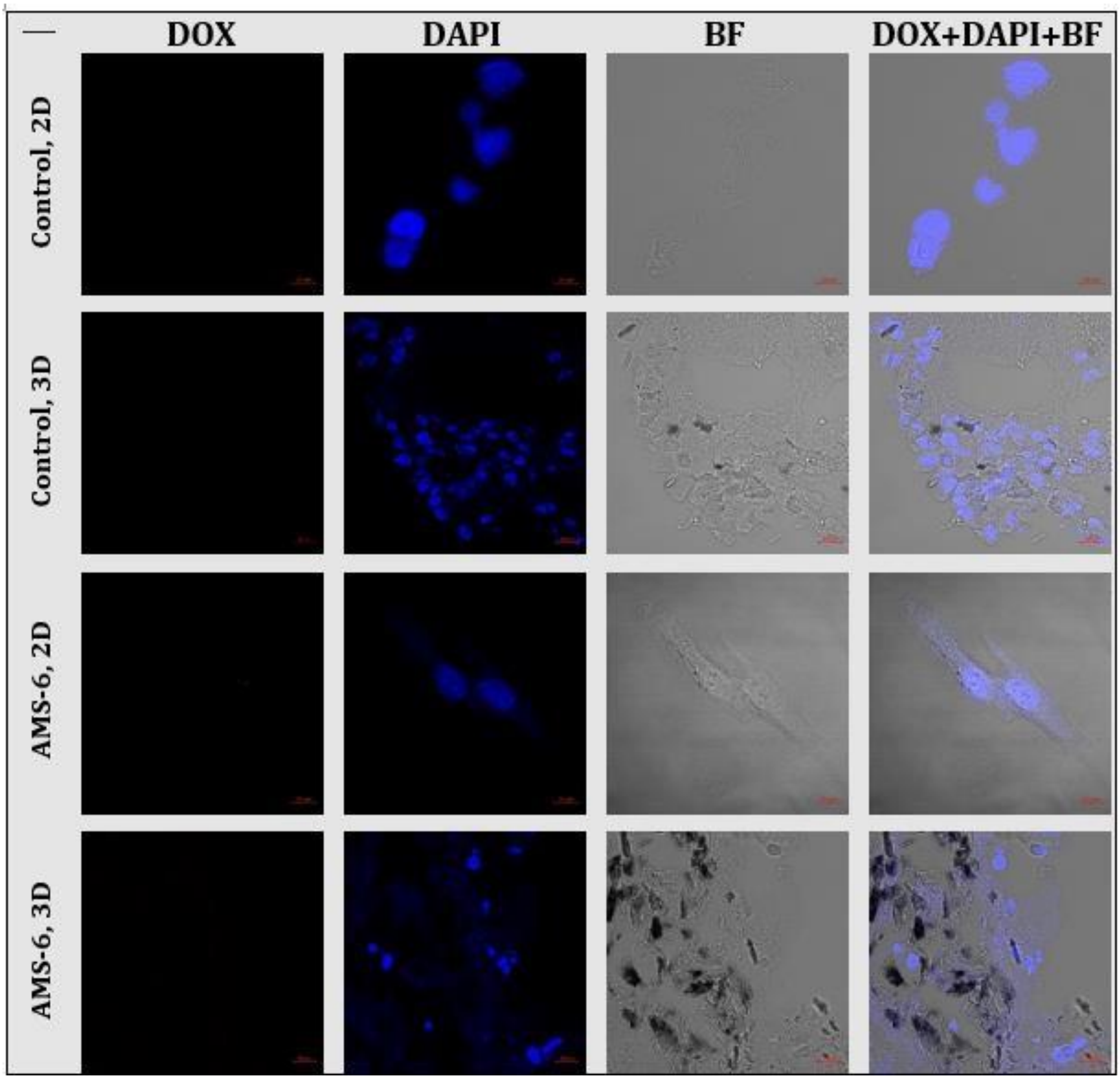


**Figure S15. Laser-scanning confocal fluorescence microscopy images 2D and 3D TECs in vitro cultures of MDA-MB-231 cells incubated with complete culture medium (control 2D and 3D) and pure AMS-6 nanoparticles for 24 h.** Note the absence of Dox fluorescence in the red channel (Dox), the clear association of DAPI fluorescence staining with cell nuclei (blue channel, DAPI). Control bright-field (BF) images were taken for visualization of the tissue structures, and an overlay of DOX, DAPI and BF show colocalization of the Dox and DAPI fluorescence. AMS-6 concentration, 50 µg/mL.

### ANGIOGENIC ASSAY ON CHICK EMBRYO CAM

#### ***Egg preparation and grafting procedure for angiogenic analysis***

The experiment was approved by the Animal Ethics Committee of Macquarie University (ARA 2015/006; 2015/006-2). The fertilized White Leghorn chicken eggs were incubated for 72 h, until embryonic day 3 (ED3). Then the eggs were placed in a horizontal position and left in the incubator for the next 30 min for repositioning of the embryos. Afterwards, the tops of the blunt ends of the shells were wiped with 70% ethanol, and 3-4 mL of the egg white was extracted from the bottom part of the egg by puncturing of the blunt end of the shell with a syringe needle (18G) at the angle of ~45° (Figure S16(a)). The albumin extraction resulted in decrease of the total volume of the egg and dropping of the chorioallantoic membrane (CAM), which were necessary for the further grafting procedure. After the extraction the stab holes were sealed with a sticky tape and the eggs were returned to the incubator until ED8. On ED8 the egg shells were cut on the blunt end to create a lid and expose the CAM. Following this, the grafting procedure was performed.

All the embryos were divided into 4 groups, with 10 eggs per a group. The group 1 was used as a control (labelled “Control”) and left ungrafted in order to evaluate the parameters of natural angiogenesis occurred in chick embryos during the period between ED8 and ED12. In other groups the following materials were aseptically implanted onto CAM under sterile conditions: 1) chick embryo liver AOSS soaked in complete culture media following the pre-seeding protocol (labelled “Scaffold”) for 24 h; 2) cell suspension of MDA-MB-231 cells, 2×105 cells in 60 µL of complete culture media (labelled “Cells”) and 3) the TECs, prepared as described above and cultured in vitro for 12 days prior grafting (labelled “TEC”). This time period of preliminary culture of TECs in vitro was chosen to obtain the engineered tumor samples containing the similar number of cells (~ 2×105 cells per sample) in the compared TECs and cell suspension xenografts. This period was also preferred to obtain the TECs with actively grown cell populations. The workflow of the grafting procedure is shown in Figure S16.

Following this, the images of the CAM were taken with a stereomicroscope as described in the section “Imaging of CAM in vivo” below. Then the eggs were sealed and returned back to the incubator and maintained under standard conditions with excluded rotation. Afterwards, on ED12, the eggshells were re-opened, and the imaging was repeated under the same conditions as on ED8. After the imaging session on ED12 the chick embryos were euthanized by quick decapitation. The CAM with grafter materials and control samples of ungrafted CAM were dissected, washed in PBS and studied by histological methods.

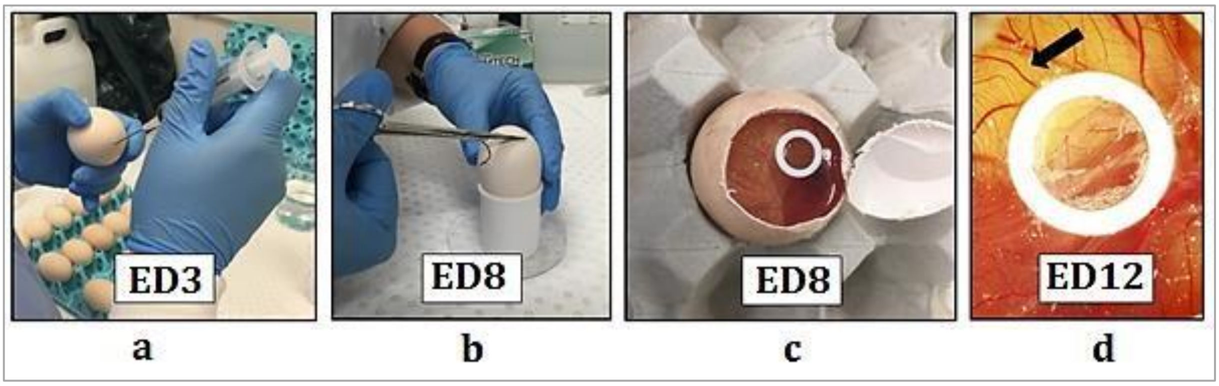


**Figure S16. Summary of grafting protocol.** (a) Extraction of the egg white. (b) Opening of the egg. (c) Implantation on the CAM. The cell suspension grafting is shown; a sterile white rubber ring taken from a 1 mL cryovial is used to prevent the leakage of the suspension. (d) The image of the same egg on ED12. The arrow shows a bifurcation of a blood vessel as a preferential grafting site.

#### ***Imaging of CAM in vivo***

The CAM of the eggs with open shell lids were imaged in vivo using the Olympus MVX10 (Olympus, Japan) stereomicroscope equipped with an eyepiece magnification ranging between 0.63× to 6.3×, a fixed focus and two objective lenses, 1×/0.25 N.A. and 2×/0.5 N.A. The microscope featured long working distance (20 to 87 mm), adjustable field-of-view (FOV, 1.7 to 55mm in diameter). In order to maintain the healthy state of the chick embryos the heating stage ThermoPlate by Tokai Hit (Shizuoka-ken, Japan) was used during in vivo inspection of CAM. The objective 1×/0.25 N.A. was applied for the rough focusing and the lens magnification of 2× was used for the detailed imaging. The samples were illuminated from above by a 100 W mercury lamp. In order to achieve a higher contrast of the red colour of the blood, a filter cube with blue illumination light was used. This filter cube contained one single-edge short pass dichroic beam splitter (Semrock, USA) with transmission of the wavelengths above 750 nm and a single-band bandpass (510570 nm) filter as the emission filter by (Semrock, USA). The microscope was coupled with a motorized Z focus ProZ Stand (Prior Scientific, USA), which provided seamless zooming from 40× to 1250× magnification (used for the focusing and study purposes).

#### ***Quantification of angiogenesis***

We evaluated the morphological parameters of angiogenesis by determining quantitative features of newly developed vessels within CAMs on ED8 and ED12. We analysed images of five eggs for each experimental group (intact eggs, cell suspensions, scaffolds, and TECs). In particular, we quantified the density (number per unit area) and length of major blood vessel sections between the branching points (segments), the blood vessels of the first order (extremities) [5] and the density of branching points [6] (Figure S17). In this approach a branch from one major vessel creates two new vessels; also the end of a vessel is counted as a branching point. Other angiogenesis parameters are calculated as described elsewhere [5] and [6].

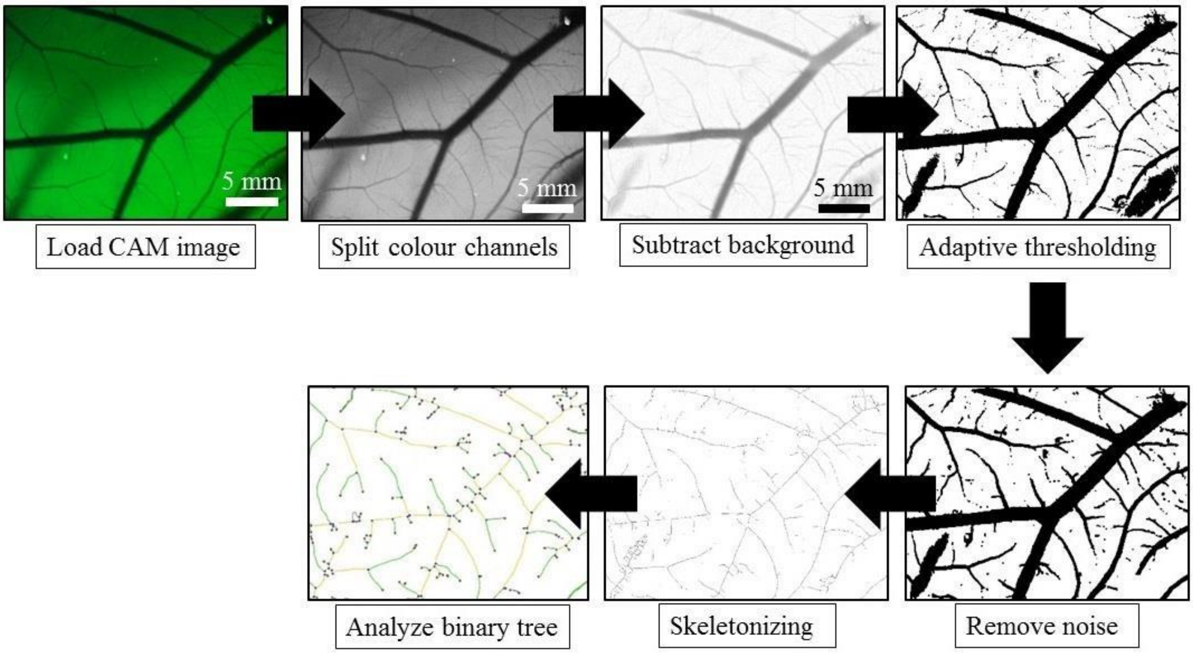


**Figure S17. Algorithm used for the validation of induced blood vessel growth.** Angiogenesis Analyzer was applied to skeletonized images. Red dots in the final panel represent the branching points whereas the green lines represent the vessels of first order (extremities) and the yellow lines show the vessels of second order as defined in [5].

The main steps of the image analysis algorithm are the subtraction of the background of the image, the adaptive thresholding, noise reduction and the skeletonization as described in References [7] and [5]. To reduce the noise erosion and dilation transformation were applied individually on each image.

After skeletonizing, the ImageJ plugin Angiogenesis Analyzer [8] was applied to each egg, four ROIs were selected, representing four opposite sides of the graft. The plugin estimates the number of branching points and total vessel length, whereas the vessels are divided into different elements. The detailed description and illustration of terminology used for quantification of binary tree of blood vessels structure can be found in the ImageJ documentation [8]. Statistical significance of the differences between the studied groups' parameters of blood vessel trees was analysed with use of one sample Kruskal-Wallace test, followed by comparisons in the pairs of the groups with use of U-test by Mann-Whitney.

The results (Figure S18) show a statistically significant increase of the following parameters per unit area in TECs on ED12 in comparison to intact controls (normal embryonic angiogenesis, ED12): the total blood vessel length, or vascular density ($p=0.07$) (Figure S18 (a)); the number of junctions ($p=0.020$) (Figure S18 (b)); c) the length of segments (the parts of the blood vessels tree between enclosed 2 neighbour branching points) ($p=0.003$) (Figure S18 (c)); the number of first order vessels (extremities) ($p=0.014$) (Figure S18 (d)); the number of branches ($p=0.039$) (Figure S18 (e)); and statistically non-significant increase in the length of branches ($p=0.121$). Statistically significant increase of the number of branching points per area ($p=0.020$) is illustrated in the main text of the paper (see Figure 6 (g). Unseeded scaffolds and cancer cell suspensions were found not to affect normal embryonic angiogenesis as there are no statistically significant differences between the scaffolds and intact controls.

There were no statistically significant differences in the parameters of the blood vessel tree architecture measured on ED12 between Scaffolds and Intact CAM, Cells and Intact CAM, and between Scaffolds and Cells.

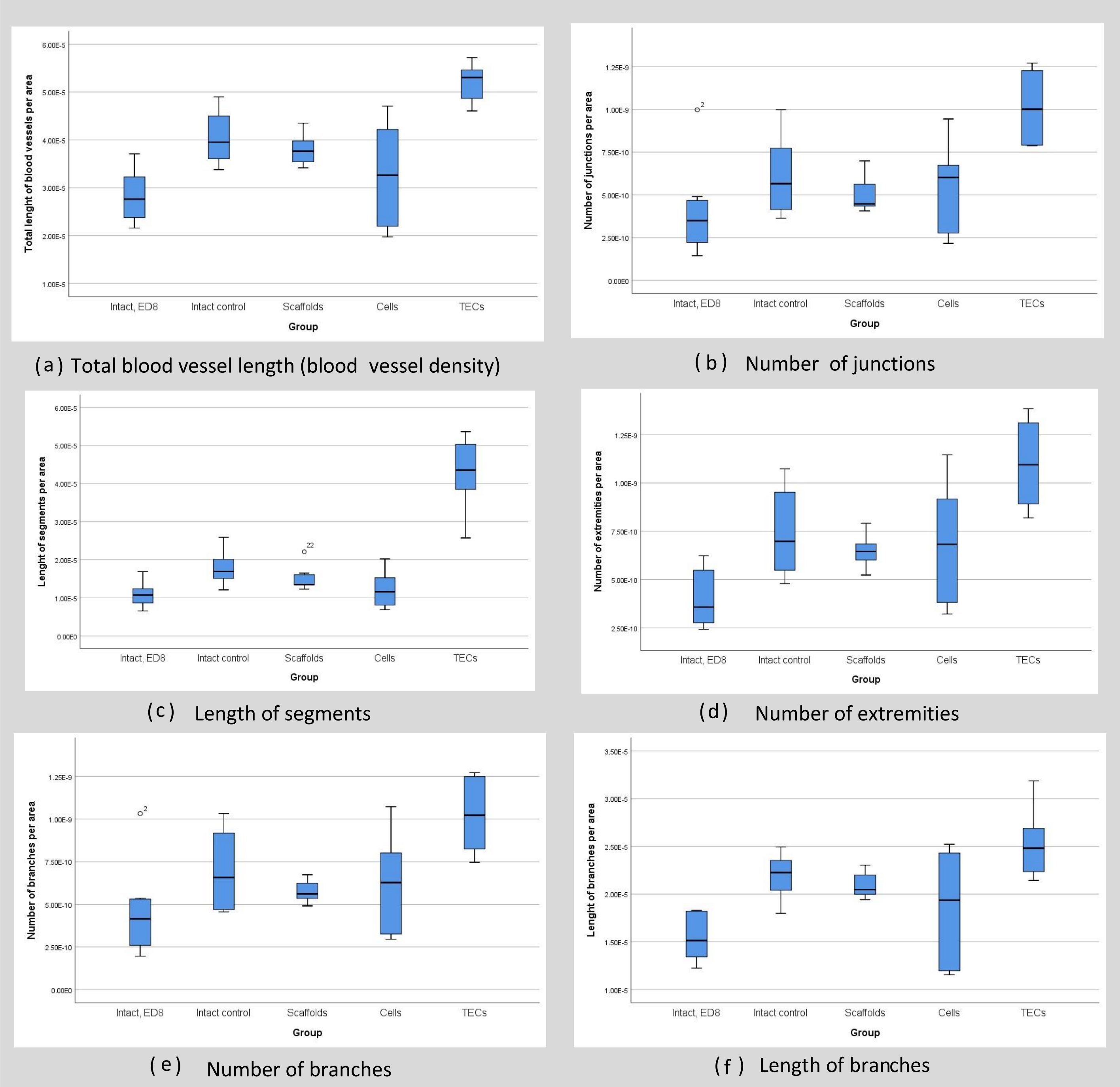


**Figure S18. Quantitative analysis of angiogenesis.** The parameters of the blood vessel tree are reflected as values per unit area. Note that increase of total blood vessel density after grafting of TECs (a) can be mainly attributed both to the intensive branching of blood vessels (b) and their increased torturing (c). The results are presented as boxplots, where thick dark horizontal line shows mean values, blue boxes indicate $CI_{95\%}$ for mean and the whiskers indicate minimal and maximal observed values

**Table S6. Testing of the statistical hypotheses for the differences between the studied groups, p- values.**

| Compared distributions | p-value | Conclusion |
|---|---|---|
| **Intact Control, ED12 vs. Intact control, ED8** | $2.6\times10^{-6}$ | Statistically significant difference |
| **Scaffolds vs. Intact control, ED12** | $1.4\times10^{-3}$ | Statistically significant difference |
| **TECs vs. Intact control, ED12** | $6.7\times10^{-2}$ | Not statistically significant difference |
| **TECs vs. Scaffolds** | $1.2\times10^{-10}$ | Statistically significant difference |
| **TECs vs. Cells** | $1.4\times10^{-4}$ | Statistically significant difference |

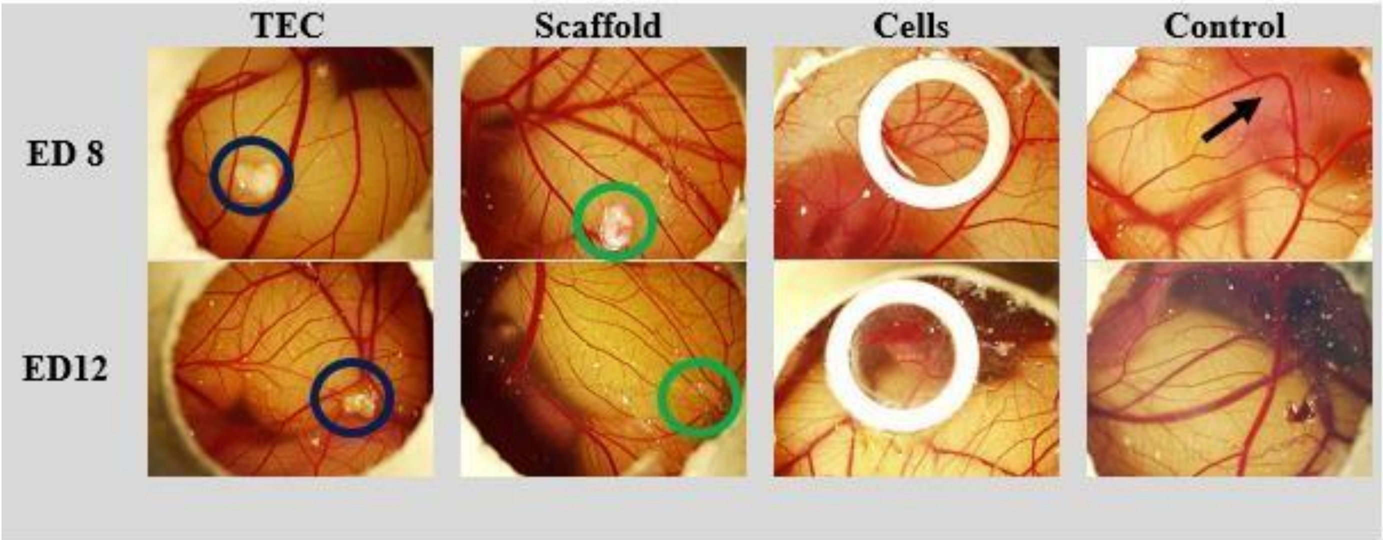


Figure S19. Architecture of CAM blood vessels in studied groups. Low-magnification (×0.63) view of control and grafted CAM on ED 8 and ED12 obtained using bright-field microscopy. Angiogenic effects induced by the following chick embryo CAM grafts: 3D *TEC* (TEC, labelled with blue circles), liver AOSSes (*Scaffold*; labelled with green circles), and suspension of MDAMB-231 cells (*Cells*; xenograft suspension was placed into the chamber formed by a white rubber ring), in comparison with natural embryonic development of CAM vasculature (*Control*) during a period between ED8 and ED12. Note that the degree of vascularization of CAM on ED8 was naturally variable between embryos. Representative images of the same eggs on ED8 and ED12 were used for analysis and shown for each group. Equal volume samples of the grafted TECs and AOSSes were used. The number of cells ($\sim2\times10^5$) in the cellular xenograft and TECs were approximately equal. An arrow in Control CAM, ED8, indicates the chick embryo residing underneath the CAM. White dust on the surface of CAM is small fragments of the eggshell dropped on the CAM during the shell windowing. Note blood vessels converging towards the grafts in *TECs* and *Scaffolds* groups.
